# Spatially resolved transcriptomic and proteomic profiling reveals cell interaction programs that predict Barrett’s esophagus progression

**DOI:** 10.64898/2026.05.08.723546

**Authors:** Isis Diaz Monarrez, Eun Na Kim, Krio Moon, Ann-Marie Baker, Phyllis Zixuan Chen, Dario Bressan, Ahmad Miremadi, Massimiliano di Pietro, Gregory J Hannon, Trevor A Graham, Rebecca C. Fitzgerald, Young Hwan Chang, Lizhe Zhuang

**Affiliations:** Department of Biomedical Engineering and Computational Biology Program, Oregon Health and Science University, Portland, OR, USA; Department of Pathology, Seoul National University Hospital, Seoul National University College of Medicine, Seoul, Republic of Korea; Centre for Evolution and Cancer, Institute of Cancer Research, London, United Kingdom; Early Cancer Institute, Department of Oncology, University of Cambridge, Cambridge, UK; Cancer Research UK Cambridge Institute, Li Ka Shing Centre, University of Cambridge, Cambridge, UK; Knight Cancer Institute, Oregon Health and Science University, Portland, OR, USA

## Abstract

Barrett’s esophagus (BE) is the precursor lesion of esophageal adenocarcinoma (EAC). It affects approximately 5% of adults in the United States and significantly increases the risk of developing EAC. However, current surveillance strategies cannot reliably distinguish patients who will progress from those who will remain stable. Direct studies of progressor BE are extremely limited due to availability of tissue with known progression outcomes, and have largely been restricted to genomic profiling approaches. The premalignant cellular landscape of progressor BE remains poorly understood.

Here, we used complementary spatial transcriptomic and proteomic imaging to profile 34 non-dysplastic BE patients under endoscopic surveillance, including those who subsequently progressed to dysplasia or EAC, termed “Progressors” and those who remained stable, termed “Non-progressors”. Transcriptomics based Xenium analysis captured 974,604 cells across 70 whole-biopsy regions, while protein based imaging mass cytometry profiled 372,242 cells across 119 selected regions. FUME-TCRseq further quantified T cell clonotypes from matched tissues scrolls. Cellular composition was generally similar between Progressors and Non-progressors. However, Progressors showed increased intestinal Barrett’s columnar cells, B cells and gastric progenitor-like cells, together with enhanced immune-epithelial interactions, whereas Non-progressors retained coordinated stromal organization.

Spatial interaction features strongly outperformed cell composition and density for progression prediction. Combined spatial interaction model achieved an area under the curve (AUC) of 0.97, compared with 0.62 and 0.68 for comparison and density alone. Complementary imaging mass cytometry further resolved the underlying immune programs, identifying cytotoxic and antigen presenting myeloid features enriched in progressors, and CD56⁺ associated memory T cell interactions enriched in non progressors.

Together, these findings support a model that BE progression is driven by progressive remodeling of epithelial-immune-stromal architecture rather than emergence of distinct dysplasia-like cell subsets. Increased T cell clonal diversity and recruitment of cytotoxic and antigen-presenting immune niches may also reflect an evolving response to genomic alteration prior to dysplasia. These results establish spatial tissue architecture, rather than specific cell types, captures progression associated microenvironmental states in BE and provides a framework for spatially informed patient stratification and early cancer risk assessment.

## Introduction

Barrett’s esophagus (BE) is a premalignant condition that affects approximately 5% of adults in the United States^1^. BE is characterized by the replacement of normal squamous epithelium with columnar epithelium exhibiting intestinal and gastric features, typically arising in response to chronic tissue injury such as gastroesophageal reflux^2,3^. BE represents the only established precursor lesion for esophageal adenocarcinoma (EAC)^2^, thus its identification is clinically important given its associated risk of progression to EAC. However, BE remains substantially underdiagnosed, with estimates suggesting that up to 80% of affected individuals have not received a diagnosis^4^. Although BE confers an increased risk of malignant transformation, the absolute annual risk of progression is low, and most patients remain clinically indolent, with progression risk estimates varying widely across studies depending on cohort composition and follow-up duration^1,3,4^. This biological and clinical heterogeneity has contributed to ongoing debate regarding the relationship between BE and EAC development, including whether a subset of EAC may arise through alternative pathways lacking a detectable metaplastic precursor. However, recent integrated epidemiologic and molecular analyses^5^ demonstrated that BE-positive and BE-negative EAC share remarkably similar genomic and transcriptional features, supporting intestinal metaplasia as a central component of EAC pathogenesis regardless of clinically detectable BE status. Together, these observations highlight the complexity of BE progression and underscore the need for improved approaches to identify progression-associated biological states and risk stratification markers.

Considerable effort has focused on identifying the genomic features in BE prior to neoplastic progression^6^. Prior studies have reported alterations in epithelial differentiation, immune infiltration, and stromal remodeling associated with disease state in cancer^7,8^. Increasing evidence indicates that the spatial organization of cells within the tissue microenvironment plays a critical role in shaping disease progression in BE and EAC. Local cell–cell interactions can modulate immune surveillance, epithelial plasticity, and stromal support, influencing whether tissue remodeling maintains homeostasis or promotes malignant transformation^9,10^. Protein-based spatial imaging approaches, including multiplex immunohistochemistry (mIHC)^11^ and CO-Detection by indEXing (CODEX)^12^, have enabled detailed characterization of the immune, stromal, and epithelial architecture of BE and EAC^13,14^. These studies have shown that progression is associated not only with changes in immune composition but also with reorganization of immune-stromal and immune-epithelial neighborhoods^14,15^. However, antibody-based approaches are constrained by predefined marker panels^16^, limiting their ability to capture broader transcriptional programs and diverse cellular states across the tissue microenvironment^8,17^.

Spatial transcriptomics provides a complementary framework for interrogating epithelial, stromal, and immune niches at broader molecular resolution, with larger biomarker panels than are typically achievable using protein based platforms. At the same time, protein based imaging remains an important complementary approach, particularly for resolving functional immune states, as certain immune markers are less effectively captured by transcriptomic panels. In this study, we combine transcriptome-scale spatial profiling with complementary multiplex protein imaging to identify microenvironmental features associated with BE progression. Using Xenium spatial transcriptomics, we define progression-associated spatial patterns through cell phenotyping, density analysis, spatial interaction modeling, and supervised classification. Complementary imaging mass cytometry (IMC) further resolves immune cell states underlying these spatial patterns. Together, we leveraged each modality for its complementary strengths, rather than performing direct cross-platform integration on non-co-registered sections. These analyses reveal that spatial organization of the epithelial-immune-stromal interface, rather than cell composition alone, captures progression-associated microenvironmental states in BE and provides a framework for spatially informed patient stratification.

## Results

### Cell phenotyping defines the major cellular landscape

We collected original formalin fixed paraffin embedded (FFPE) tissue blocks from 34 patients with non dysplastic BE. Among these, 17 patients subsequently progressed to dysplasia or esophageal adenocarcinoma after a mean follow up of 2.2 ± 0.9 years, whereas the remaining 17 patients have remained non progressors to date, with a mean follow up duration of 6.6 ± 1.1 years (**Fig. 1A**). For each tissue block, multiple sections and scrolls were prepared between 2022 and 2024 for IMC, Xenium spatial transcriptomics, and DNA/RNA extraction (**Fig. 1B**). Some samples were not included in the Xenium dataset because of tissue exhaustion or incomplete tissue architecture within the available sections (**Fig. 1C**).

**Figure 1.**
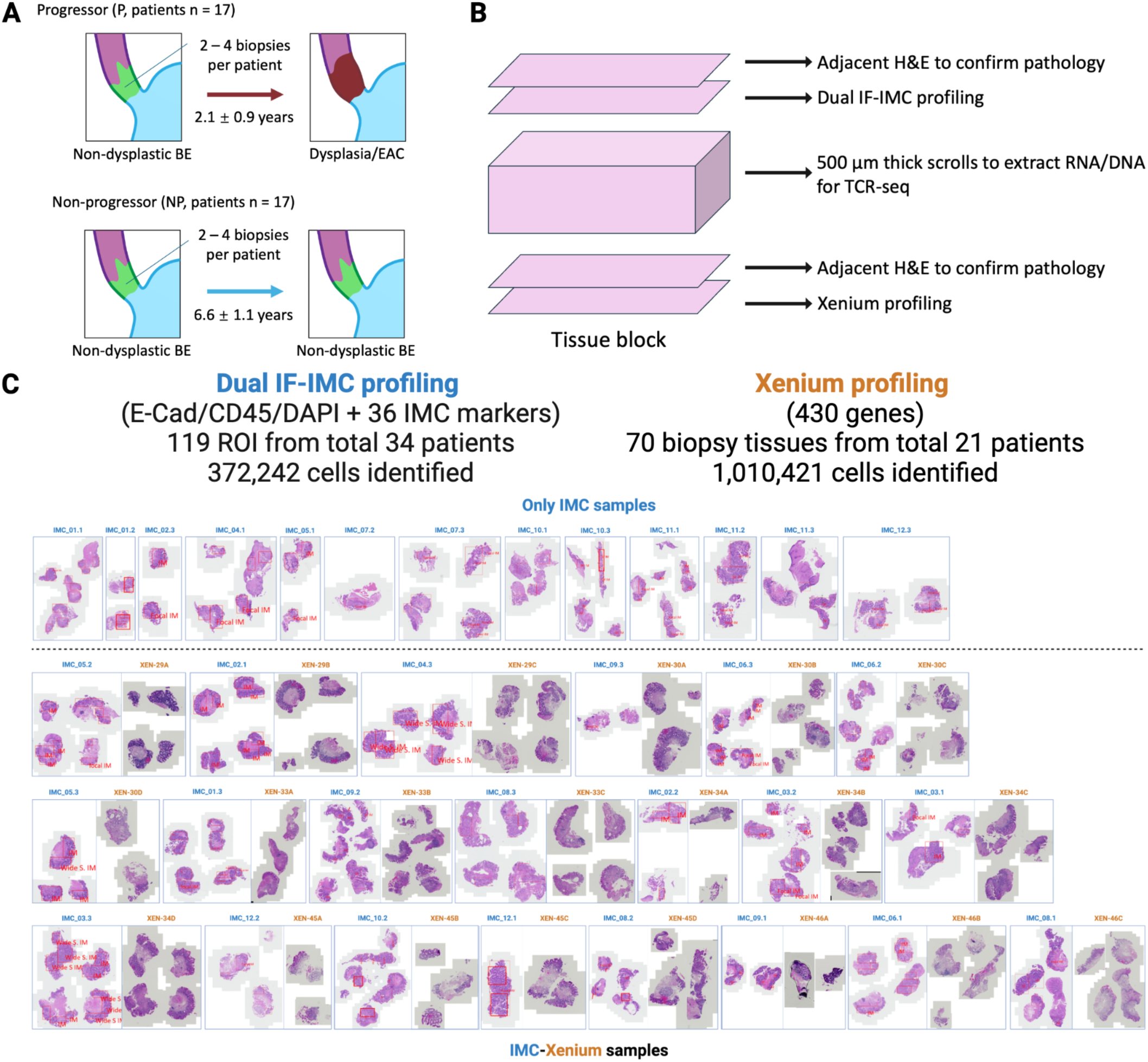
Study cohort and distinct spatial profiling designs for Xenium and IMC. **A,** Progressors and Non-progressors cohorts. **B,** Tissue block processing workflow. Blocks were initially sectioned for dual IF-IMC profiling, followed by collection of 500 µm scrolls for DNA/RNA extraction and TCR-seq. Additional sections were subsequently generated for Xenium spatial transcriptomics after the technology became available. **C,** IMC and Xenium profiling designs. IMC was performed on 119 selected rectangular ROIs, providing sampled but not full-tissue coverage. Xenium was performed on 70 biopsies with full tissue coverage. Because IMC and Xenium were performed on non-adjacent sections generated under separate study designs, the datasets were analysed as complementary, non-matched spatial profiles.

We first focused on the Xenium dataset as the primary discovery platform, leveraging its high resolution spatial transcriptomic profiling across 430 genes (**Supplementary Table 1**). In total, the Xenium dataset comprised 70 biopsy tissues, with each tissue section defined as a single region of interest (ROI). This included 41 Progressor ROIs and 29 Non-progressor ROIs, encompassing 1,024,921 cells in total. (**Fig. 2A**).

**Figure 2.**
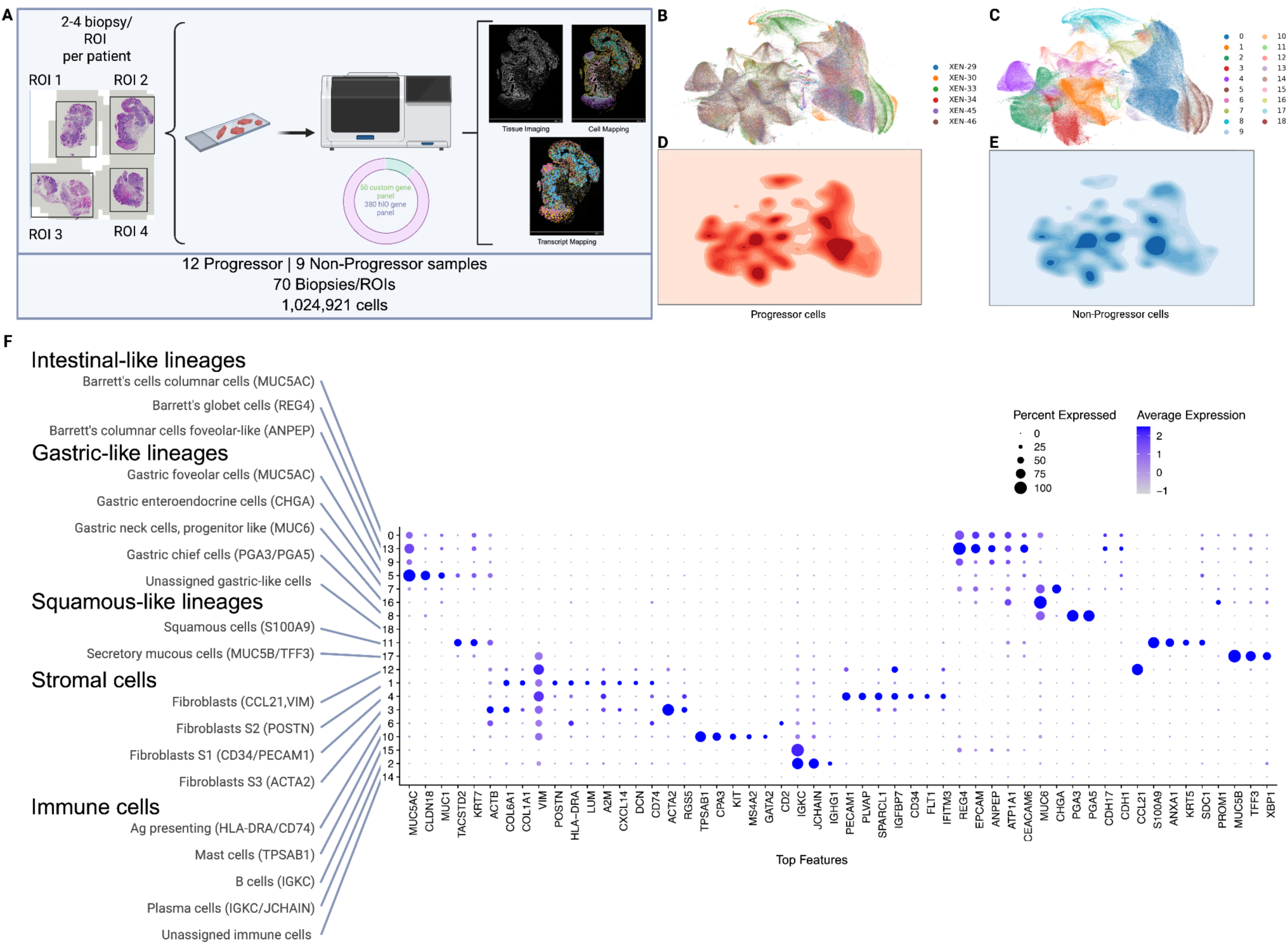
Spatial transcriptomic profiling identifies 19 cell phenotypes. **A,** Cohort design and Xenium spatial transcriptomics workflow. Biopsies from Non-progressor and Progressor patients were profiled using a 430-gene panel, yielding approximately 1 million cells across 70 biopsies. **B–C,** UMAP projections of all cells colored by patient (**B**) and cell phenotype clusters (**C**). **D–E,** Progressor and Non-progressor cells mapped onto the UMAP, highlighting differences in cell population abundance. **F,** Dot plot of marker genes used for cell phenotype annotation. Dot size indicates the percentage of cells expressing each marker, and color indicates scaled expression.

Following quality control to remove outlier cells not associated with tissue regions and potential contaminant island cells, 960,104 cells were retained for downstream analysis (**Supplementary Fig. 2A** and **Supplementary Table 2**). Evaluation of probe and gene detection metrics revealed variability in transcript detection across genes; positive probe per cell–gene analysis showed that 37 genes exhibited an average of more than three probes per cell per gene (**Supplementary Fig. 2B**). It is noteworthy that some common immune markers such as CD3 and CD8 were not effectively detected under Xenium. These observations supported the need for normalization, which was performed using the Seurat pipeline (Methods). Examination of sample-level distributions demonstrated good mixing, indicating minimal batch effects across Xenium slides (**Fig. 2B**).

Unsupervised clustering of transcriptomic profiles identified 19 transcriptionally distinct cell populations (**Fig. 2C**). To determine whether transcriptional states differed by progression status, UMAP density distributions were stratified by condition (**Fig. 2D-2E**). Overall, epithelial, stromal, and immune populations showed substantial overlap between Progressor and Non-progressor samples, with no discrete clusters or unique transcriptional states specifically enriched in either condition. These findings suggest that dysplastic progression in BE is unlikely to be primarily driven by the emergence of distinct cell populations, but may instead reflect alterations in cellular abundance, spatial organization, and tissue architecture.

The identified cell populations were grouped into five major biological categories based on marker expression: intestinal-like, gastric-like, and squamous-like epithelial lineages, as well as stromal and immune populations (**Fig. 2F**). The Xenium dataset captured heterogeneous epithelial populations, including both gastric and intestinal lineages, consistent with previous reports^18,19^. Cell lineages were annotated using established BE associated markers, including REG4^+^/OLFM4^+^ goblet and progenitor cells^20^, MUC5AC^+^/MUC6^+^ gastric-like epithelium^21^, and KRT5^+^/S100A9^+^ squamous-like cells. Stromal clusters were characterized by COL1A1^+^/POSTN^+^/VIM^+^ fibroblast populations, while immune populations were identified using marker profiles consistent with those reported in prior BE spatial analyses^12^ (**Fig. 2F**).

### Cell Density reveals progression-associated enrichment patterns

Because cell composition alone did not clearly distinguish progressor from non-progressor tissues, we next examined cell density metrics that account for variability in tissue area across ROIs. To focus on reproducible patterns, we applied a 25% patient-level prevalence threshold, retaining only clusters detected in at least 25% of patients for both density and composition analyses (Methods).

ROI level hierarchical clustering of cell density patterns revealed progression associated enrichment patterns that were not apparent from compositional analysis alone (**Fig. 3A** and **Supplementary Fig. 3A**). To quantify these differences, we summarized mean cell density per ROI and compared distributions between progression groups. This analysis identified several populations with significant differential enrichment. Non-progressor tissues showed increased densities of Fibroblast S2 cells, while Progressor samples were enriched for Barrett’s columnar cells (cluster 0), IGKC⁺ B cells (cluster 15), and MUC6⁺/CHGA⁺ Gastric neck like progenitor cells (cluster 16) (**Fig. 3B**).

**Figure 3.**
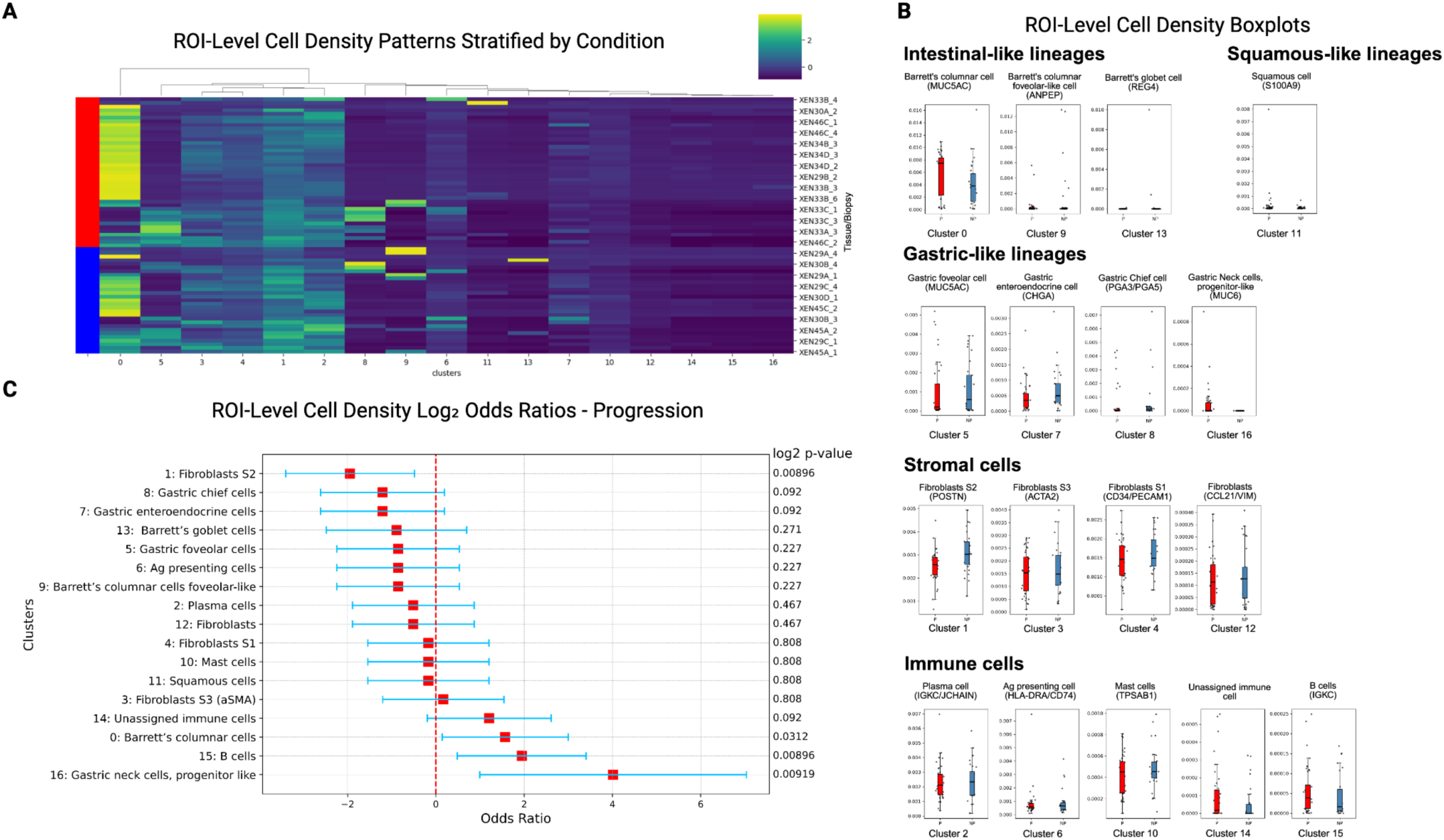
ROI-level cell density patterns show limited separation by progression status. **A,** Heatmap of cell densities across ROIs, stratified by progression status: Progressor (red) and Non-progressor (blue). **B,** Forest plot showing log₂ odds ratios for progression, highlighting cell populations enriched or depleted in progressor ROIs. **C,** Boxplots showing ROI-level cell densities across annotated cell populations.

Log-odds ratio analysis further quantified these associations using both composition-based and density-based approaches (**Fig. 2C**; Methods). Positive log2 odds ratios indicate preferential enrichment in progressor tissues, whereas negative values indicate enrichment in non progressors. Density-based odds ratios identified multiple populations with significant condition-associated enrichment. Fibroblast S2 cells (cluster 1) were preferentially enriched in Non-progressors, whereas B cells (cluster 15), Gastric neck progenitor-like cells (cluster 16), and Barrett’s columnar cells (cluster 0) were preferentially associated with Progressor tissues (**Fig. 3C**). In contrast, composition-based analysis identified only two clusters with significant progression associated enrichment, and these appeared to be driven by a small number of outlier samples rather than consistent enrichment across the cohort (**Supplementary Fig. 3B-3C**). This further supports the notion that alterations in local cell abundance, rather than overall proportional composition, better distinguish progression associated tissue states.

To further contextualize composition-based findings, we examined cell composition at the broader lineage level by aggregating clusters into immune, stromal, and epithelial compartments. Condition-level heatmaps did not reveal strong global compositional differences between Progressor and Non-progressor samples (**Supplementary Fig. 4A**). However, composition-based boxplots and odds ratio analyses consistently identified Fibroblast S2 (cluster 1) and Gastric enteroendocrine cells (cluster 7) as associated with Non-progression, whereas B cells (cluster 15) and Gastric neck progenitor-like cells (cluster 16) were associated with Progression (**Supplementary Fig. 4B and 4C**). These observations support the consistency of the composition-based analyses, although the magnitude of these differences remained modest and insufficient to provide robust classification between conditions.

**Figure 4.**
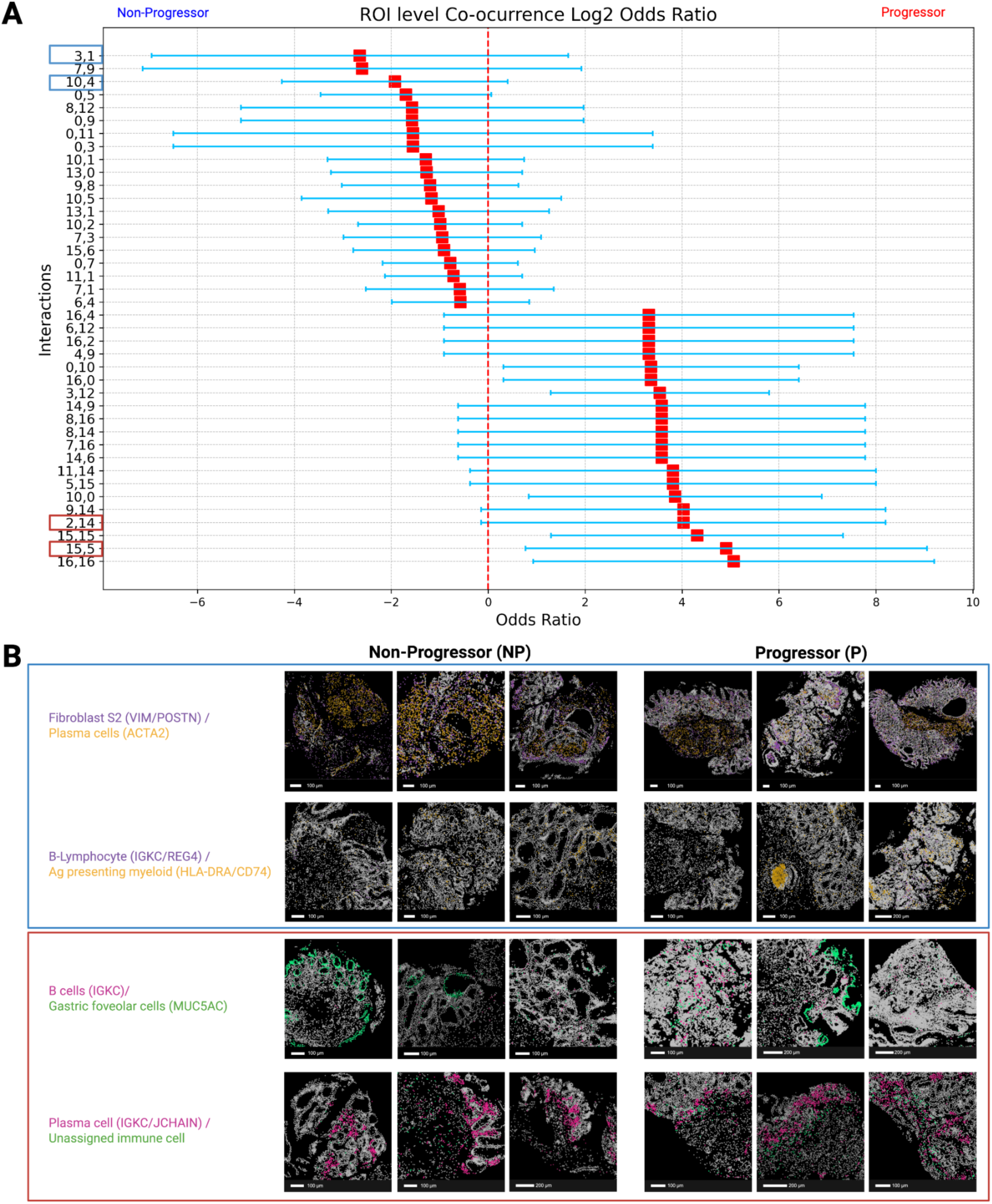
Spatial co-occurrence patterns differ by progression status. **A,** Forest plot of log₂ odds ratios for ROI-level cell-cell co-occurrence, highlighting cell pairs enriched or depleted in Progressor ROIs. **B,** Representative spatial maps showing selected interactions in Non-progressor (blue) and Progressor (red) ROIs, illustrating progression-associated differences in cell proximity patterns.

Together, these findings suggest that increased density of specific cell types, rather than composition alone, better reflects progression-associated tissue remodeling in non-dysplastic BE^9,10^. However, density differences alone do not capture how cells are organized relative to one another, motivating examination of cell–cell interaction patterns.

### Spatial cell-cell interaction patterns distinguish progressor and non-progressor tissue microenvironments

Building on density-associated enrichment patterns, we next quantified cell–cell interactions to determine whether tissue organization differed between progression groups beyond cell abundance. We applied two complementary approaches: co-occurrence analysis, which assesses whether two cell types tend to appear together within the same local region; and neighborhood enrichment, which evaluates whether cell types are enriched within spatial neighborhoods^13,22^. Co-occurrence captures larger scale spatial co-localization patterns, whereas neighborhood enrichment reflects more short range cell–cell adjacency, together providing complementary views of spatial tissue organization at different spatial scales (**Supplementary Fig. 5**). Both analyses were performed with GraphCompass normalization^14^ (Methods) to minimize inter-ROI variability, and log-odds ratios were calculated to identify interaction cell type pairs preferentially associated with Progressor or Non-progressor.

Progressor tissues showed enriched co-occurrence patterns consistent with increased immune-epithelial spatial co-localization and immune niche remodeling (**Fig. 4A** and **Supplementary Fig. 6A**). Spatial co-localization between B cells and MUC5AC⁺ Gastric foveolar-type epithelium (cluster pair 15 and 5) suggests co-localized immune-epithelial neighborhoods within metaplastic regions. This observation is consistent with prior spatial studies in BE reporting structured immune niches enriched for B-lineage cells, supporting a role for adaptive immune engagement in dysplastic-prone epithelium contexts^23^. Co-occurrence of plasma cells with unassigned immune populations (cluster pair 2 and 14) further supports altered immune organization in Progressors, in line with previous reports showing plasma cell co-localization with myeloid and CD4⁺ T-cell populations and progression-associated restructuring of immune niches^12,24^. Representative examples of these interactions are shown in **Fig. 4B**. Notably, neighborhood enrichment analysis also identified mast cells-Barrett’s columnar cells interactions (cluster pairs 10 and 0) and Gastric chief cells-Unassigned immune cells (cluster pair 8 and 14) , supporting the consistency of selected spatial patterns across complementary metrics (**Supplementary Fig. 6B**).

In contrast, Non-progressor tissues exhibited interactions consistent with preserved stromal organization and structured immune-stromal positioning (**Fig. 4A** and **4B**). Spatial co-localization between ACTA2⁺ Fibroblast S3 and POSTN⁺/HLA-DRA–enriched Fibroblast S2 populations (cluster pair 3 and 1) suggests coordinated stromal organization involving contractile and matrix-remodeling fibroblast states^25^. Enrichment of Mast cells-Fibroblast S1 neighborhoods (cluster pair 10 and 4) further indicated maintained immune-stromal positioning in Non-progressor tissues, potentially supported by canonical stromal niche signaling and mediator-driven fibroblast activation^26,27^. These stromal interaction patterns were also observed by neighborhood enrichment analysis, including Fibroblast S2-Mast cell (cluster pair 1 and 10) and Gastric-like epithelium-Mast cell (cluster pair 5 and 10) adjacency (**Supplementary Fig. 6B**).

Collectively, these findings show that differences between Progressor and Non-progressor tissues extend beyond cell abundance to distinct patterns of spatial organization. Progressor tissues were characterized by increased immune-epithelial proximity and immune niche remodeling, whereas Non-progressor tissues maintained more coordinated stromal and immune-stromal organization. Importantly, several interaction pairs were consistently identified by both co-occurrence and neighborhood enrichment analyses, supporting the robustness of these spatial patterns and prompted us to evaluate whether integrating these spatial features could improve prediction of progression status.

### Spatial interaction features improve supervised classification of progression status

To evaluate whether the spatial features identified above provide additional predictive value for distinguishing between Progressor and Non-progressor tissues, we applied a supervised binary classification framework using support vector machine (SVM) classifiers with leave-one-out cross-validation (LOOCV) (Methods).

Models trained on composition or density features alone showed limited predictive capacity, yielding AUC values of 0.62 and 0.68, respectively (**Fig. 5A**). Incorporating spatial interaction features improved classification performance, with neighborhood enrichment features increasing the AUC to 0.81 and co-occurrence features achieving the highest performance (AUC = 0.97). These results indicate that spatial interaction structure captures progression-associated information not reflected by cell abundance alone.

**Figure 5.**
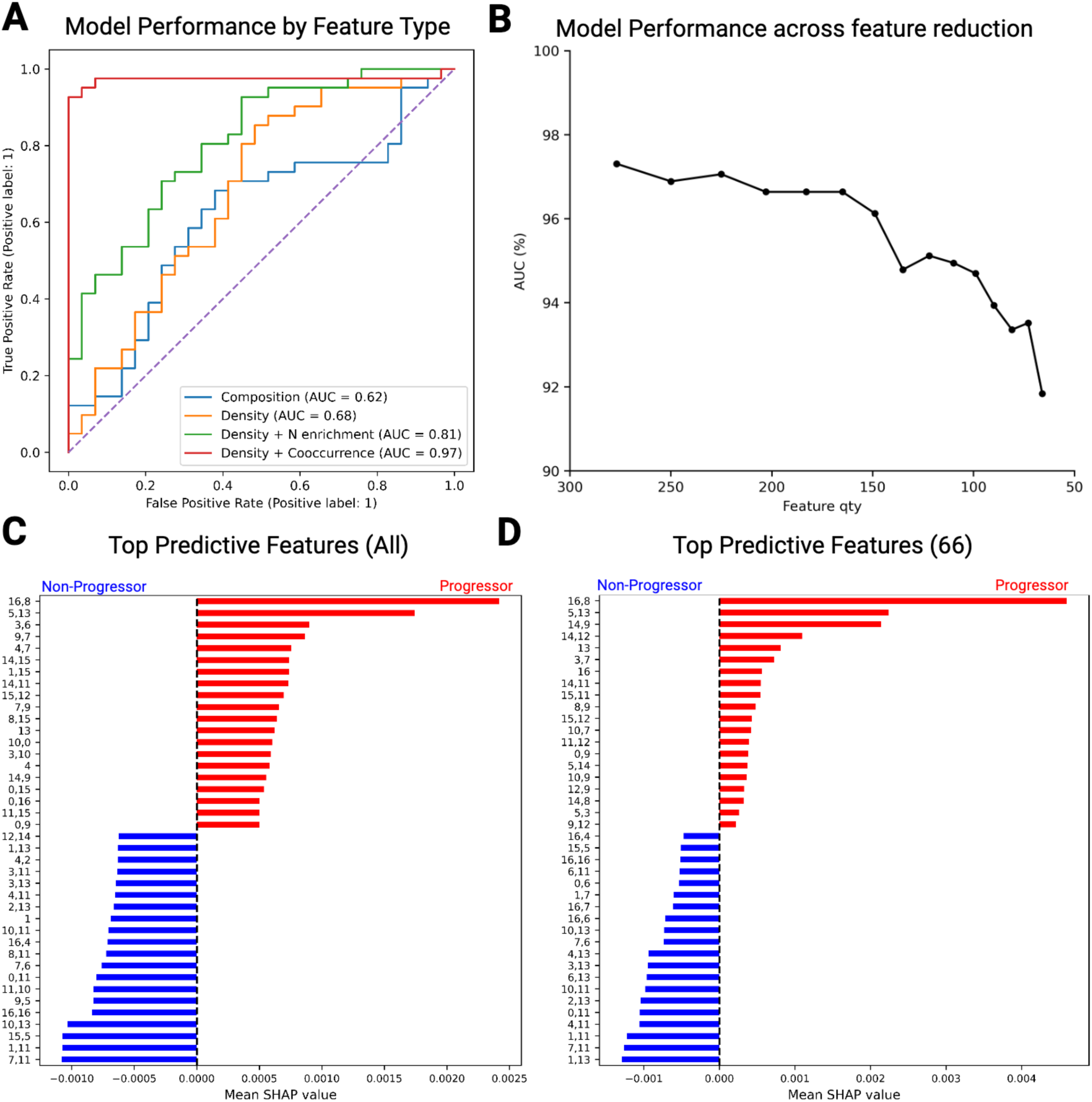
Predictive modeling identifies progression-associated spatial interactions. **A,** ROC curves comparing models trained using cell composition, cell density, neighborhood enrichment, and cell-cell co-occurrence features. **B,** AUC across feature reduction, showing model performance stability. **C–D,** SHAP-based feature importance before (**C**) and after (**D**) feature reduction. Top-ranked features highlight spatial interactions most strongly associated with progression.

To assess model stability and identify a compact predictive feature set, we perform iterative feature reduction guided by SHapley Additive exPlanations (SHAP)^15^ importance score, removing the lowest-contributing 10% of features at each step (**Fig. 5B**). Classification performance remained robust across feature-reduction thresholds, with peak performance observed using the full feature set and a gradual decline as features were removed. A reduced model retaining 66 spatial features maintained strong discriminatory performance, indicating that a core subset of spatial interactions carries much of the classification signal.

SHAP-based feature importance analysis identified biologically coherent interactions among the strongest contributors to model performance (**Fig. 5C**). Not all interactions with significant odds ratios were prioritized by the classifier, highlighting the distinction between univariate enrichment and multivariate predictive contribution. Among Progressor-associated features, Gastric neck cell-Gastric chief cell (cluster pair 16 and 8), Gastric foveolar cell-Barrett’s goblet cell (cluster pair 5 and 13), and Fibroblast S3-Antigen-presenting myeloid cell (cluster pair 3 and 6) interactions ranked highly, suggesting a mosaic gastric-intestinal lineage state, and immune-stromal remodeling in Progressor tissues. Non-progressor-associated features included Gastric enteroendocrine cell-Squamous-like cell (cluster pair 7 and 11) and Fibroblast S2-Squamous cell (cluster pair 1 and 11) co-occurrence, suggesting retention of squamous associated epithelial programs, relative absence of intestinal lineages, and a less remodeled stromal microenvironment in Non-progressor tissues.

Several key interactions, including Gastric neck cell-Gastric chief cell (cluster pair 16 and 8) and Gastric foveolar cell-Barrett’s goblet cell (cluster pair 5 and 13) co-occurrence, remained among the top predictors for Progressors in the reduced 66-feature model (**Fig. 5C** and **5D**). Together, these findings show that spatial interaction features substantially improve prediction of progression-associated tissue states, and that coordinated cell-cell interaction programs provide a core classification signal not captured by composition or density alone. Given that certain immune markers, including CD3 and CD8, and stromal markers such as COL1, were less effectively detected at the mRNA level in the Xenium dataset, we incorporated the complementary IMC dataset to provide protein level characterization of immune and stromal states and interactions.

### IMC profiling resolves immune states associated with progression-linked spatial programs

As outlined in **Fig. 1B**, the tissues used for IMC were sectioned prior to those used for Xenium, but were derived from the same cohort. IMC and Xenium were performed on sections from non-adjacent tissue depths (**Fig. 1B**) and differed in spatial coverage, with Xenium ROIs representing whole biopsy tissue IMC ROIs representing selected rectangle regions within a biopsy tissue (**Fig. 1C** and **Supplementary Fig. 1**), therefore, direct cross-platform integration at single-cell or region-level resolution was not performed. Instead, each modality was leveraged for its complementary strengths: Xenium provided transcriptome-scale spatial discovery across larger tissue areas, whereas IMC enabled protein-based characterization of immune, epithelial and stromal phenotypes within selected ROIs (Methods; **Fig. 6A** and **6B**).

**Figure 6.**
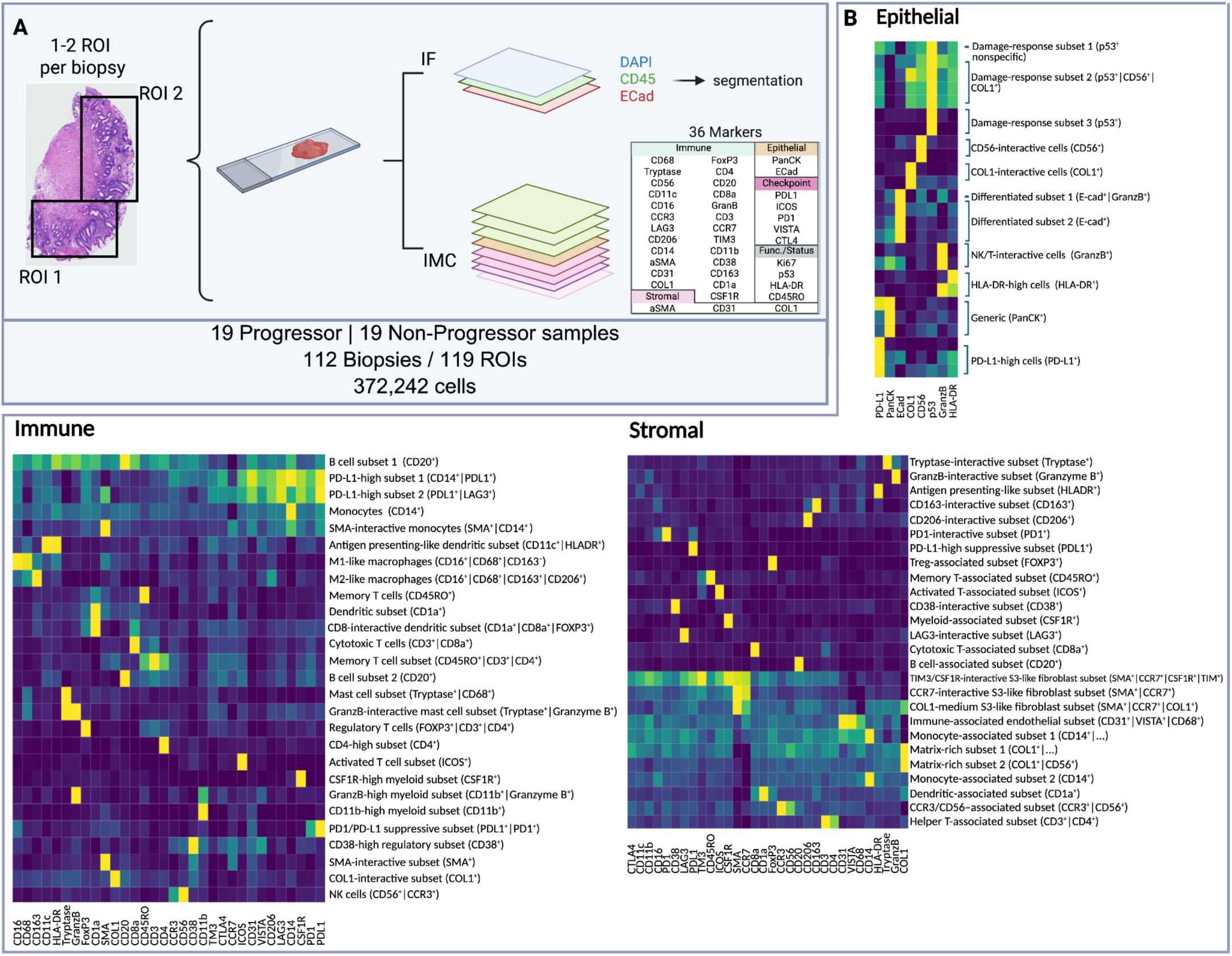
IMC profiling identifies three major cellular compartments comprising 64 cell phenotypes. **A,** IMC cohort design and workflow. Non-progressor and Progressor biopsies were profiled using a 36-marker panel, yielding ∼350,000 cells across 119 ROIs. **B,** Cell phenotyping heatmap of epithelial, immune, and stromal cell populations.

We applied the dual IF-IMC approach^11^ using IF (immunofluorescent) CD45 and ECad as master lineage markers alongside 36 IMC markers (**Supplementary Table 1**) across 119 selected ROIs spanning Progressor and Non-progressor biopsies, identifying 372,242 cells in total (**Fig. 6A**). Cells were first categorized as immune (IF CD45⁺), epithelial (IF ECad⁺), or stromal (IF CD45⁻/IF ECad⁻), and were then clustered independently. The phenotyping identified diverse cell populations, especially immune and stromal subsets (**Fig. 6B**). Consistent with Xenium analysis, cell proportions alone showed only limited progression-associated differences, although odds ratio analysis identified selected populations enriched in Progressor or Non-progressor tissues (**Fig. 7A**). These results further supported that progression associated tissue states are not defined solely by differences in cell abundance, but also by altered spatial and microenvironmental organization.

**Figure 7.**
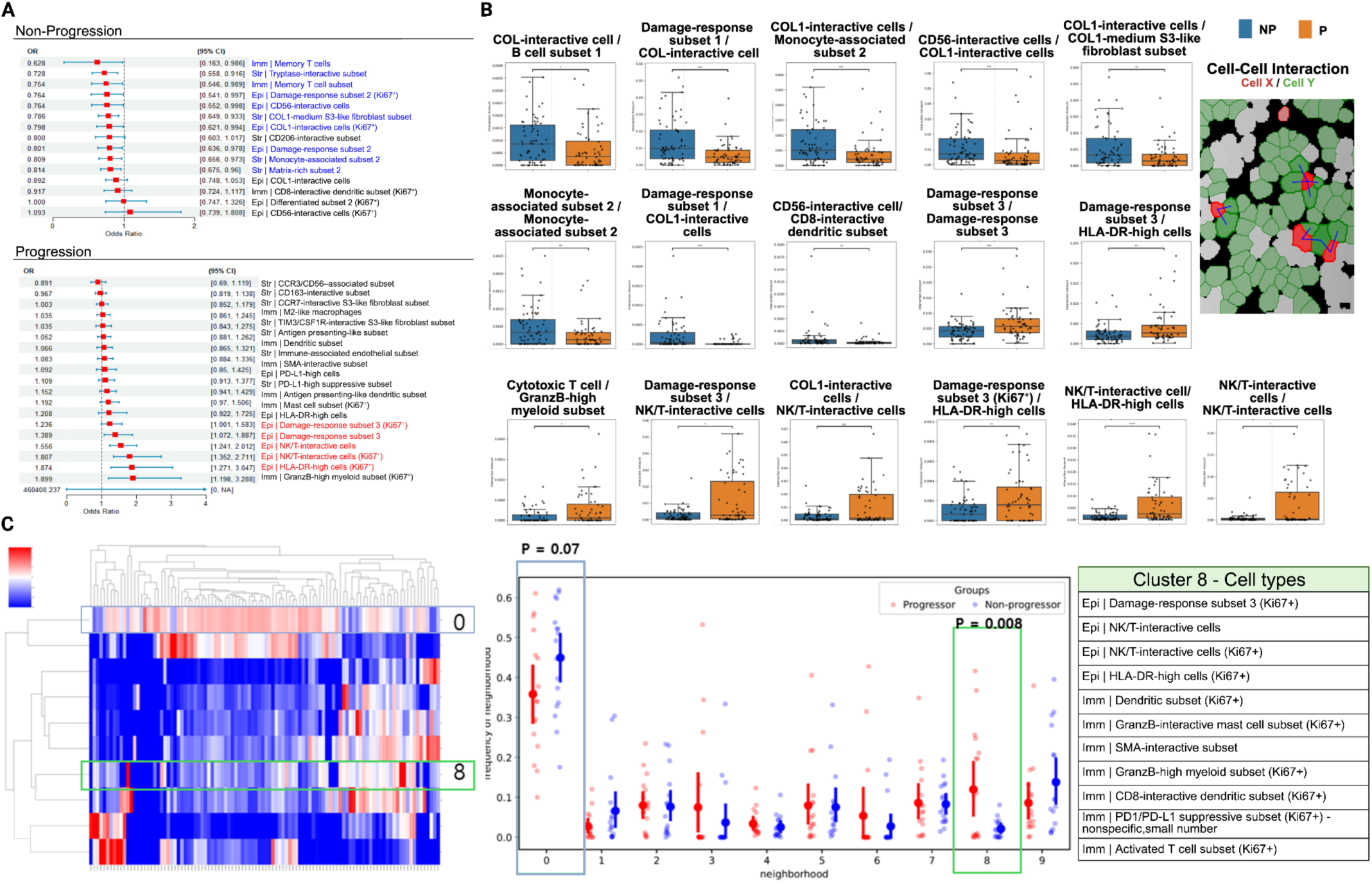
IMC identifies progression-associated cell–cell interactions and predictive spatial features. **A,** Odds ratios of cell proportions associated with progression status. **B,** Cell–cell interaction analysis highlighting significant interactions between annotated cell populations. **C,** i-niche analysis identifying neighborhood 8 as progression-associated proliferative immune–epithelial niche enriched for damage-response epithelial cells, antigen-presentation-associated epithelial states, dendritic/myeloid populations, and activated T-cell subsets.

We next examined IMC-derived cell-cell interactions to determine whether protein-defined spatial organization differed by progression status (**Fig. 7B**). Progressor tissues showed increased interactions involving epithelial-associated immune activation markers and cytotoxic or antigen-presenting immune populations. In particular, Granzyme B^+^ and HLA-DR^+^ epithelial-associated regions showed increased proximity to CD11b⁺Granzyme B⁺ myeloid cells and CD11c⁺HLA-DR⁺ dendritic cells, consistent with an immune-activated and remodeled microenvironment in Progressor tissues (**Fig. 7B** and **Supplementary Fig. 7**). Representative spatial maps illustrated these progression-associated interaction patterns within selected ROIs (**Supplementary Fig. 7**).

In contrast, Non-progressor tissues showed spatial patterns consistent with more preserved epithelial-immune and immune-stromal organization. IMC revealed enrichment of CD56⁺ cell-epithelial interactions and collagen-associated epithelial regions, together with increased CD45RO⁺CD3⁺CD4⁺ memory T-cell populations. These features suggest a tissue microenvironment characterized by maintained epithelial architecture and structured immune positioning in Non-progressors (**Fig. 7B** and **Supplementary Fig. 8**).

To identify the most predictive IMC-derived interaction features, we applied Random Forest analysis to the cell-cell interaction profiles. This analysis highlighted interaction features associated with each condition, including immune-activated and antigen-presenting programs in Progressor tissues and CD56⁺-associated or memory T-cell-associated interactions in Non-progressors (**Supplementary Fig. 8**). In addition, protein-based i-niche analysis^28^, which defines local cellular neighborhoods based on the composition of cells immediately surrounding an index cell , identified neighborhood 8 as progression-associated (**Fig. 7C**). This neighborhood was composed of multiple proliferative epithelial and immune cell states, including Ki67⁺ damage-response epithelial cells, NK/T-interactive epithelial cells, Ki67⁺ HLA-DR-high epithelial cells, dendritic cell subsets, Granzyme B-high myeloid cells, activated T cells, and CD8-interactive dendritic subsets. These features suggest that neighborhood 8 captures a higher-order immune-epithelial niche characterized by epithelial stress/proliferation, antigen-presentation-associated epithelial states, and local immune activation. A small PD1/PD-L1 suppressive subset was also present, although this population was limited in number and should be interpreted cautiously. Together, the enrichment of neighborhood 8 in Progressor tissues supports the presence of a recurrent immune-remodeled epithelial microenvironment associated with BE progression.

Although IMC did not directly recapitulate the exact Xenium-defined interaction pairs, both platforms supported a shared biological theme: Progressor tissues showed increased immune-epithelial interactions and immune associated remodeling, whereas Non-progressor tissues retained more structured epithelial, stromal, and immune organization. IMC further refined these observations by resolving protein-based immune phenotypes and spatial interaction features associated with these broader programs. Together, these orthogonal spatial profiling approaches indicate that BE progression is associated with coordinated remodeling of immune-epithelial and immune-stromal organization, rather than changes in cell abundance alone.

### FUME-TCRseq reveals increased T-cell clonal diversity in Barrett’s Progressor tissues

To further characterize the adaptive immune microenvironment, we performed FUME-TCRseq^29^ on RNA extracted from tissue scrolls from the same FFPE blocks (**Fig. 1B**). FUME-TCRseq profiled the expressed TCRβ CDR3 region, enabling individual T cell clones to be identified and quantified. We first assessed whether each sample possessed a distinct T cell repertoire using Morisita index, which showed very low overlap between samples (**Fig. 8A**). This is anticipated as the samples were derived from different patients and BE is known to bear highly heterogeneous mutations^30^.

**Figure 8.**
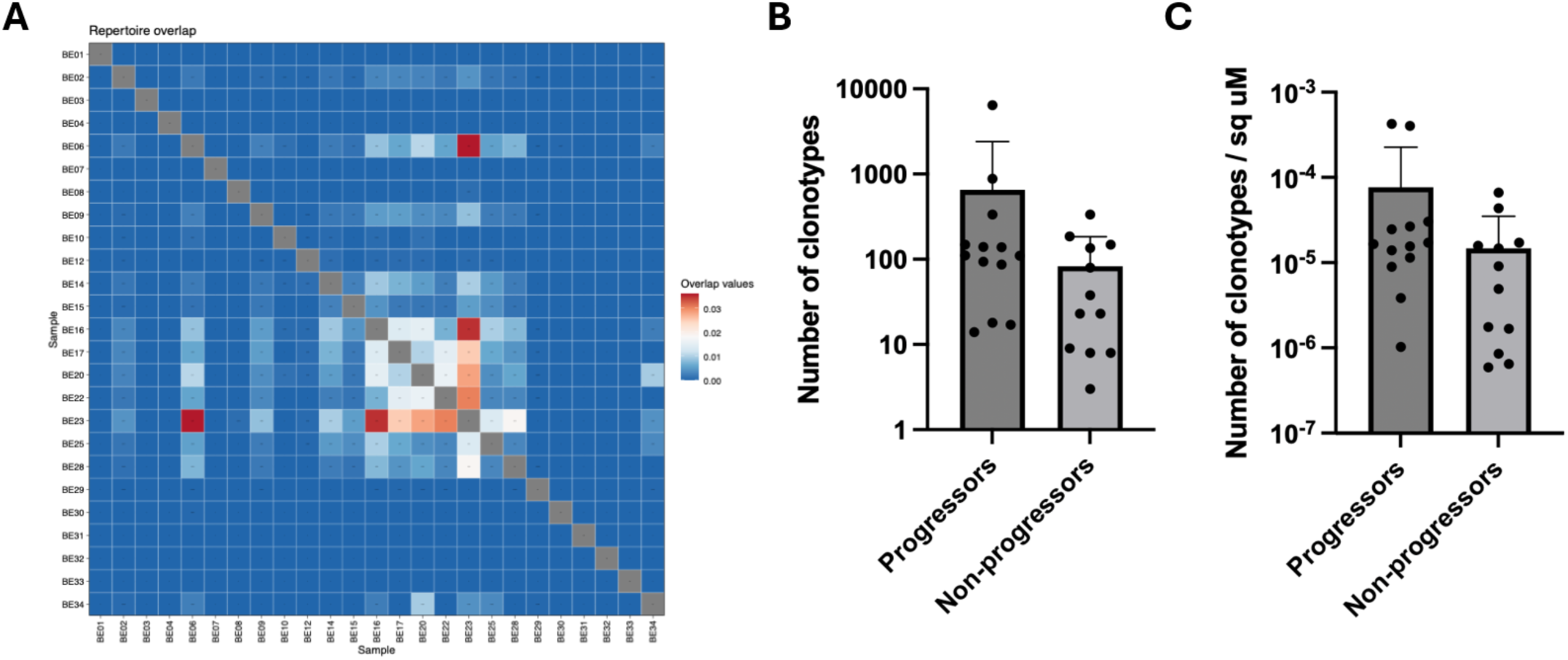
FUME-TCRseq reveals increased T cell clonal diversity. **A**. Morisita overlap between each sample. **B**, Raw counts of the number of unique clonotypes in the TCRseq data. **C**, Clonotypes count normalized to tissue areas.

We then quantified the T cell clonal abundance. Progressor tissues showed a higher number of unique T-cell clonotypes than Non-progressor tissues, both in raw clonotype counts and after normalization to tissue area (**Fig. 8B** and **8C**). These findings suggest that the immune niche remodeling observed in Progressors in the Xenium and IMC analyses is accompanied by a broader and more diverse infiltration of T cell clones. This increased clonal diversity in Progressors aligns with the IMC findings of enriched cytotoxic and antigen-presenting myeloid features. This may indicate active and ongoing immune recruitment against heterogeneous epithelial alterations during BE progression, consistent with the observation that progressors harbor higher levels of copy number alterations years prior to progression^6^.

## Discussion

Risk stratification and prediction in non dysplastic BE has been a major focus of research in early cancer detection. Current management of Barrett’s oesophagus relies on resource intensive endoscopic surveillance, despite the fact that the majority of patients remain indolent, limiting the overall efficiency of the system. However, direct research in this area has been greatly hindered by the limited availability of appropriate tissue samples, which require specimens collected years prior to neoplastic progression, namely from “progressors” in our study. Only a handful of studies have assembled such cohorts, but these are typically limited to only tens of cases^6,12,31,32^, whereas the majority of the literature has focused on precancerous tissues collected at or after the emergence of dysplasia or cancer.

Prior studies of BE progression have primarily focused on sequencing-based genomic profiling, whereas spatial characterization of the BE tissue microenvironment remains comparatively limited. Earlier spatial work identified immune alterations accompanying the transition from non-dysplastic BE to dysplasia, including changes in T-cell abundance within the epithelial compartment^33^. Subsequent mIHC studies demonstrated progressive remodeling of immune cell composition across BE, dysplasia, and esophageal adenocarcinoma, implicating spatial immune architecture in disease progression^34^. These observations additionally motivated clinically oriented spatial pathology approaches such as TissueCypher, which integrates multiplex biomarker imaging with tissue architecture features for BE risk stratification and has demonstrated improved predictive performance compared with standard clinicopathologic variables^35^. More recently, high-plex CODEX profiling extended these findings by demonstrating that progression is associated not only with compositional changes, but also with reorganization of immune and stromal neighborhoods, including spatial interactions involving plasma cells and stromal–immune compartments^12^. Despite these advances, current protein-based and biomarker-driven spatial approaches remain constrained by predefined marker panel molecular depth, and challenges associated with tissue heterogeneity and sampling variability^16,36,37^ In particular, recent evaluations of BE biomarker assays and surveillance strategies have emphasized limitations related to cohort size, prospective validation, and the impact of spatial heterogeneity on risk prediction performance^37,38^. Here, we applied Xenium spatial transcriptomics profiling to provide a broader characterization of progression-associated microenvironmental programs and spatial cellular interactions in BE.

Our analyses indicate that spatial organization of epithelial-immune-stromal interface, rather than cellular composition alone, is a central feature distinguishing Progressor from Non-progressor BE tissues. Initial compositional analyses showed only modest differences between groups, consistent with prior spatial protein-based studies^12^. However, incorporating spatially informed features, including cell density and cell-cell interaction patterns, revealed progression-associated microenvironmental programs that were not fully resolved by abundance-based measures. These findings support a model in which tissue architecture encodes clinically relevant information beyond the presence or proportion of individual cell types.

Density-based analysis identified condition-associated enrichments that were not evident from composition alone. Fibroblast S2 populations were preferentially enriched in Non-progressor tissues, whereas B cells and Gastric neck progenitor-like cells were increased in Progressor lesions. This distinction suggests that progression-associated remodeling in BE may be reflected by density-based enrichment of specific epithelial and immune states, rather than by proportional redistribution alone. Because density metrics incorporate tissue area and absolute cell number, they may be better suited than compositional measures to capture localized changes in tissue organization. These findings are broadly consistent with prior observations linking stromal organization with Non-progressor states and increased immune activation with progression^12^.

Spatial interaction analysis further revealed distinct microenvironmental programs associated with progression status. Progressor tissues exhibited increased immune-epithelial proximity, including spatial coupling between B cells and Gastric foveolar-type epithelium and co-occurrence of Plasma cells with immune populations, consistent with immune niche remodeling within metaplastic regions. These patterns align with prior IMC and CODEX studies reporting reconstructing of immune neighborhoods, particularly involving plasma cell and myeloid compartments, in progressing BE tissues^11^. In contrast, Non-progressor tissues demonstrated coordinated stromal organization, including Fibroblast S3-Fibroblast S2 proximity and Mast cells-Fibroblast S1 adjacency, consistent with maintained stromal architecture. Together, these findings suggest that stable BE lesions retain more organized stromal and immune-stromal structure, whereas progression is associated with immune-centered remodeling of the epithelial interface (**Fig. 7B**).

A notable feature of our findings is the consistent spatial patterns observed across transcriptomic and protein-based platforms. The increased immune-epithelial proximity observed in Progressor tissues by Xenium was supported by IMC, which revealed enrichment of cytotoxic and Antigen-presenting immune populations, including Granzyme B⁺ myeloid cells, HLA-DR⁺ dendritic cells and M2-like macrophages, in proximity to epithelial compartments. Conversely, Non-progressor tissues showed preservation of CD56⁺-associated and memory T-cell-associated interaction. Stromal-myeloid spatial interactions also emerged as a recurrent feature: [fibroblast S3]-[antigen-presenting myeloid cell] co-occurrence was identified as a top feature in the Xenium-based classification model, and HLA-DR⁺ myeloid populations were similarly associated with progression-linked spatial patterns in IMC analysis (**Fig. 7C**). Prior CODEX profiling has also reported enrichment of myeloid-stromal niches in progression-associated contexts^12^, suggesting that stromal-immune interfaces may represent a conserved microenvironmental axis in BE progression. The convergence of these observations across orthogonal spatial modalities strengthens their biological plausibility and supports further evaluation as candidate spatial biomarkers.

Supervised classification analyses further underscored the relevance of spatial organization. The improvement in cross-validated discriminatory performance from composition and density features to spatial interaction features suggests that cell-cell organization carries progression-associated information not captured by abundance metrics alone. Notably, interactions identified as statistically enriched by odds ratio analysis were not uniformly prioritized by predictive modeling, highlighting the distinction between univariate enrichment and multivariate predictive contribution. Some enriched interactions may carry redundant information, whereas others with modest individual effects may contribute more strongly when integrated with complementary spatial signals. Feature reduction analyses further suggested that a core subset of interactions, including epithelial lineage coupling and fibroblast-myeloid interfaces, retained substantial discriminatory signals, reinforcing their relevance as components of progression-associated spatial architecture.

Clinical decision making in Barrett’s oesophagus is still largely driven by the emergence of dysplasia. Emerging image based technologies are increasingly moving the field towards AI powered analyses that link histopathological images with specific cellular states and altered genomic features^40–43^. Our findings support the idea that disrupted tissue architecture may represent an important but underexplored component that could be integrated into future image based risk stratification approaches, ideally using longitudinal samples.

The strengths of this study are threefold. First, we analysed a true progressor cohort of non dysplastic Barrett’s oesophagus cases that subsequently developed dysplasia. Second, we applied complementary spatial profiling approaches spanning both transcriptomic and protein based platforms, together with TCR sequencing, generating a valuable multidimensional dataset. Third, the use of whole tissue analysis rather than tissue microarrays maximised tissue representation and increased the capacity to identify spatially relevant features^44–46^. Collectively, this study provides a conceptual framework for future spatial analyses in cancer risk stratification and a valuable resource for further data driven investigation.

Several limitations should be noted. Our analyses were based on a limited number of patients, ideally, a larger longitudinal cohort will be required to fully capture inter-patient heterogeneity and validate classification performance. Because supervised models were evaluated using LOOCV in a small cohort, the AUC values should be interpreted as evidence of cross-validated discriminatory signal rather than as established clinical prediction performance. Additionally, while spatial associations provide insight into microenvironmental organization, they do not establish causality; functional studies will be required to determine whether the identified interaction programs actively contribute to progression or reflect secondary tissue remodeling. Xenium and IMC analyses were performed on non-adjacent tissue sections and differed in tissue coverage, precluding direct cross-platform alignment at single-cell or region-level resolution. IMC was therefore used for complementary immune characterization rather than direct multimodal integration. Future studies incorporating co-registered multimodal profiling, larger independent validation cohorts, and orthogonal functional assays will be important to extend and mechanistically interpret the spatial programs identified here.

In summary, this study shows that spatial organization of the epithelial-immune-stromal interface captures progression-associated microenvironmental programs that are not apparent from cellular composition alone. By combining transcriptome-scale spatial profiling, protein-level imaging, and supervised classification, we identify convergent spatial features that distinguish progressor from non-progressor tissues across complementary analytical platforms. These findings establish spatial cell-cell interaction patterns as a promising dimension for improved risk stratification in Barrett’s esophagus and provide a framework for spatially informed biomarker development in premalignant disease.

## Resource Availability Lead contact

Requests for further information and resources should be directed to and will be fulfilled by the lead contacts, Dr. Young Hwan Chang and Dr. Lizhe Zhuang.

## Data and code availability

● The datasets presented in this article are not readily available. Requests to access the dataset should be directed to Lead contact.
● Any additional information needed to reanalyze the data reported in this paper is available from the lead author by request.

## Author Contributions

I.D.M.: conceptualization, data curation, formal analysis, investigation, methodology, project administration, software, validation, visualization, writing – original draft, and writing – review and editing. E.K.: methodology, formal analysis, investigation, methodology, project administration, software, validation, visualization, writing – original draft, and writing – review and editing. K.M.: data curation, formal analysis, and investigation. A.M.B.: FUME-TCRseq and data analysis. P.Z.C.: tissue preparation, immunostaining and acquisition of IMC dataset. D.B.: support and consultation of IMC experiments. A.M.: histopathological assessment of tissues. M.D.P.: supervision of endoscopic biopsy collection and consultation. G.J.H.: support and consultation of IMC experiments. T.A.G. support and consultation of FUME-TCRseq and data analysis. R.C.F: conceptualization, supervision, investigation, writing – review & editing. Y.H.C.: conceptualization, methodology, funding acquisition, resources, supervision, project administration, and writing – review and editing. L.Z.: conceptualization, methodology, funding acquisition, resources, supervision, project administration, and writing – review and editing.

## Acknowledgments

We thank all patients who contributed to the study. We thank Dr Ania Piskorz and Ms Rachel Barnes from the Genomics Core, and Dr Joe Heffer from the Histology Core at the Cancer Research UK Cambridge Institute, for their assistance and support with the Xenium experiments. This work was supported by the International Alliance for Cancer Early Detection (ACED) funding (EDDAMC-2023/100006 ) to Y.H.C. and L.Z, and the CRUK funding (EDDPJT-May23/100046) to L.Z. Y.H.C. also acknowledges funding from National Institutes of Health (R01 CA253860). The research reported in this publication used computational infrastructure supported by the Office of Research Infrastructure Programs, Office of the Director, of the National Institutes of Health under award number S10OD034224. Schematics were created using BioRender (https://BioRender.com).

## Methods

### Study cohort and distinct spatial profiling designs for Xenium and IMC

We analyzed tissue specimens from patients with non-dysplastic Barrett’s esophagus (BE), including Progressors and Non-progressors , using two complementary spatial profiling approaches: Xenium in situ spatial transcriptomics and imaging mass cytometry (IMC) (**Fig. 1A**). These datasets were generated using distinct sampling strategies and were therefore not treated as spatially matched measurements.

Tissue blocks were initially processed for the original IMC study design, with adjacent sections used for dual IMC profiling, followed by 500 µm scrolls collected for DNA and RNA extraction for TCR-seq. Additional sections were later generated for Xenium profiling after the technology became available (**Fig. 1B**). Consequently, IMC and Xenium were not performed on immediately adjacent sections and were not designed for direct cell-level or region-level co-registration.

The two platforms also differed in tissue coverage. IMC profiled 119 selected rectangular regions of interest (ROIs), which sampled portions of the tissue but did not cover the full biopsy area. In contrast, Xenium profiled 70 biopsies with full tissue coverage, enabling broader assessment of cellular composition and spatial organization (**Fig. 1C**, **Supplementary Fig. 1**). Given these differences in section origin, ROI selection, and tissue coverage, we analyzed Xenium and IMC as complementary but non-matched datasets, with Xenium serving as the primary discovery platform and IMC providing orthogonal protein-level characterization of immune states.

### Patient cohorts and samples

Samples were obtained via endoscopic biopsies from patients with known BE under clinical surveillance. Ethical approval was granted by the UK Health Research Authority (Biomarker REC 01/149), and all patients provided written informed consent prior to sampling. The full study cohort used for IMC comprised 34 patients, included 17 Progressors, defined as patients who progressed from non-dysplastic BE to high-grade or low-grade dysplasia (HGD/LGD) or intramucosal carcinoma with a minimum follow-up of one year. For the Progressor group, biopsies from the final endoscopy prior to the diagnosis of progression were analyzed. The control group consisted of 17 Non-progressors who maintained non-dysplastic BE throughout surveillance. Non-progressor samples analyzed here were selected from endoscopic biopsies with at least five years of confirmed clinical stability.

A subset of this cohort was profiled by Xenium spatial transcriptomics and included 21 patients total, comprising 12 Progressors and 9 Non-progressors. These samples were derived from the same clinically annotated surveillance cohort using identical progression criteria but were limited by tissue availability and assay feasibility.

For each tissue block, two initial adjacent sections were cut for histopathological assessment and dual IF-IMC profiling, respectively (see *Imaging Mass Cytometry acquisition and processing*). Subsequently, 500 μm scrolls were cut for DNA and RNA extraction for TCR sequencing. Finally, two additional adjacent sections were taken for histopathological assessment and Xenium profiling. Due to the staggered availability of these technologies, the blocks were processed in separate phases; consequently, the sections used for IF-IMC and Xenium are separated by at least 500 μm. As these sections are not adjacent, the cellular and tissue architectures vary greatly between these distinct depths within the same sample, and we did not fully integrate the IF-IMC and Xenium datasets.

### Xenium data generation and pre-processing

Tissue samples were processed using Xenium Explorer (v3.2.0). ROIs were manually defined using the Freehand Selection tool to delineate histologically relevant tissue areas. Only cells whose centroids fell within the ROI boundaries were retained for downstream analysis to ensure spatial consistency. Squamous islands, which are frequently observed in clinical biopsies taken adjacent to the normal squamous lining, were excluded from the analysis. This to ensure that the observed transcriptomic and cellular phenotypes reflect the true risk-associated divergence between Progressor and Non-progressor cohorts, rather than the inherent biological distinctions between squamous and columnar lineages. Each retained cell was assigned an ROI label in the metadata to enable ROI-level stratification in subsequent analyses. This filtering reduced the raw cell count from 1,024,921 to 974,604 cells (P = 650,009 cells; NP = 310,095 cells).

### Xenium transcriptomics profiling and data processing

Raw count matrices were analyzed in R (v4.3.3) using Seurat (v5.0.0). Low-quality cells were removed by excluding cells expressing fewer than one detected gene, resulting in a final dataset of 960,104 cells. Normalization and variance stabilization were performed using SCTransform with regularized negative binomial regression, which accounts for technical variation while preserving biological signal^47^. Principal component analysis (PCA) was performed on the scaled data, and the top 25 principal components were selected based on the elbow method. Unsupervised clustering was performed using the Louvain algorithm implemented on the FindClusters function with a resolution parameter of 0.1. Clusters were visualized using Uniform Manifold Approximation and Projection (UMAP) to facilitate interpretation of cellular heterogeneity.

### Xenium cell phenotyping and cluster annotation

Cell phenotypes were assigned using a marker-based annotation strategy. For each cluster, differentially expressed genes were identified, and a dot plot was generated to summarize marker expression patterns, displaying the top differentially expressed features based on log fold change and requiring expression in at least 25% of cells within the cluster. Cluster identities were then assigned through literature-guided annotation based on the expression of these enriched marker genes and their consistency with known cell types reported in BE and related gastrointestinal tissues.

### Xenium statistical analysis for Cluster enrichment

For downstream analyses, a 25% patient-level prevalence cutoff was applied to both composition and density metrics, excluding clusters detected in fewer than 25% of patients to reduce patient-specific bias.

To assess the association between clusters and progression status (progressor vs. non-progressor), we performed an odds ratio (OR) analyses^48^ on both composition and density using scipy.stats.contingency (SciPy v1.15.2).

For composition analysis, a “one vs all” framework was applied. For each cluster, the number of cells in progressor and non-progressor samples was compared to the combined counts of all remaining clusters across both conditions. This approach evaluates whether a given cell population is over- or under-represented relative to the overall cellular landscape.

For density analysis, clusters were binarized into high- versus low-density groups using the global median density threshold. This strategy incorporates spatial context by accounting for local cell concentration rather than absolute abundance. Odds ratios were reported as log2(OR), enabling identification of clusters enriched in progressor versus non-progressor tissues.

### Xenium spatial analysis: co-occurrence

To assess spatial co-localization patterns between cell types, we used Scimap (v2.2.11) to perform spatial interaction mapping with the radius-based method. For each cell, all neighboring cells within a 70-μm radius of its centroid were identified, and the probability of cell-type co-localization was calculated. The 70-μm radius was selected to capture immediate microenvironmental neighborhoods while minimizing inclusion of distant, potentially unrelated cells. Statistical significance was evaluated by comparing observed interactions to a permuted random background distribution, thereby estimating the likelihood of specific cell-type association beyond chance.

### Xenium spatial analysis: neighborhood enrichment analysis

To evaluate statistically significant differences in cell-type neighborhoods, we computed neighborhood enrichment for each sample using Squidpy (v1.2.3)^13^. Cell–cell adjacency graphs were constructed using Delaunay triangulation to model direct cellular contacts. The nhood_enrichment function was then used to measure the observed frequency of each cell-type pair adjacency and compare it to the expected frequency obtained through permutation testing. Enrichment values greater than one indicate over-representation, whereas values less than one indicate depletion. This analysis provides a complementary view of spatial organization that is independent of radial distance thresholds.

### Xenium spatial analysis: normalization

To account for variability introduced by tissue composition, cell-type proportions, ROI area, and patient-specific effects, we normalized both co-occurrence and neighborhood enrichment outputs using the GraphCompass framework^14^. Interactions present in ≤25% of patients were excluded prior to modeling to reduce sparsity and improve robustness yielding 150 and 261 features for co-occurrence and neighborhood respectively.

GraphCompass fits the spatial interaction values using a linear model that includes patient identity as a covariate, thereby harmonizing results across ROIs while controlling for inter-patient variability. The framework also performs *t*-tests on the normalized coefficients to assess statistical significance. This normalization reduces ROI-level confounding while preserving biologically meaningful spatial patterns for downstream analyses.

### Xenium supervised binary classification

We applied a supervised machine learning framework to evaluate whether spatial features improve stratification between Progressor and Non-progressor samples. Given the limited sample size, support vector machine (SVM) classifiers were implemented using scikit-learn (v1.7.0) with leave-one-out cross-validation (LOOCV) to maximize training data usage and minimize bias.

To assess the predictive utility of different feature sets, SVMs were trained separately using composition-based and density-based features. Density-based features consistently achieved superior classification performance and were therefore selected for subsequent modeling. Hyperparameters were optimized using grid search, yielding optimal values of C = 51, gamma = 0.001, with class_weight = balanced.

Given the large number of spatial features, we further evaluated model robustness through iterative feature reduction. Feature importance was estimated using SHapley Additive exPlanations (SHAP)^15^. At each iteration, the lowest-contributing 10% of features were removed, and the SVM was retrained using the reduced feature set. Model performance was evaluated at each step using the area under the receiver operating characteristic curve (AUC), enabling identification of an optimal balance between feature complexity and predictive performance.

### IMC acquisition and processing

IMC was performed using the Hyperion Imaging System (Fluidigm) to achieve high-dimensional spatial analysis of immune, stromal, and epithelial components in BE tissue samples.

Tissue sections (4 µm) from FFPE blocks were deparaffinized, rehydrated, and subjected to metal-conjugated antibody staining, followed by laser ablation at a resolution of 1 µm. The ablated material was subsequently analyzed via time-of-flight mass spectrometry (CyTOF) to detect and quantify multiplexed protein expression in individual cells.

IMC imaging enabled simultaneous detection of multiple markers in a single tissue section, allowing for comprehensive phenotypic characterization of immune, epithelial, and stromal cells. Image acquisition parameters were standardized to ensure consistent data quality and reproducibility across all samples.

High-resolution immunofluorescence (IF) markers, including E-cadherin, cytokeratin, and CD45, were used to enhance cell segmentation, improve feature extraction, facilitate unsupervised cell phenotyping, and enable spatial analysis with greater accuracy.

### IMC antibody panel

The panel includes markers targeting immune, stromal, epithelial, immune checkpoint, and functional statue-related markers (**Supplementary Table 1**).

Antibody Categories and Targets

- Immune markers: CD45RO, CD3, CD4, CD20, CD8a, CD1a, CD11b, CD11c, CD14,CD16, CD68, CD56, CD38, CD163
- Stromal markers: αSMA, CD31, COL1
- Epithelial markers: PanCK, E-cadherin
- Immune checkpoint markers: PD-L1, PD-1, ICOS, VISTA
- Functional status markers: Ki67, P53, HLA-DR, Granzyme B
- Additional markers: CCR3, CTLA4, CSF1R, Tryptase, CCR7, TIM3, LAG3, FOXP3

### IMC unsupervised cell phenotyping

Single-cell IMC data were clustered using GPU-accelerated Phenograph implemented with RAPIDS/CuPy-based workflows. Louvain clustering was performed with a neighborhood parameter of 40 on 372,242 cells, yielding 64 clusters (11 Epithelial | 27 Immune | 26 Stromal). Cluster identities were assigned based on z-normalized mean antibody intensity heatmaps and refined through expert manual curation by a board-certified pathologist. Final annotations were validated by overlaying cluster assignments onto raw marker images for spatial confirmation.

### IMC odd ratio calculation

To assess associations between IMC-defined cell populations and progression status, odds ratios were calculated for each cluster comparing Progressor and Non-progressor samples based on cell composition and density metrics. Statistical significance was assessed using chi-square or t-tests with a significance threshold of *p* < 0.05. Clusters showing significant differences between conditions were selected for downstream analyses.

### IMC cell-to-cell interaction analysis

To quantify spatial interactions between IMC-defined cell populations, a GPU-accelerated computational pipeline was used to identify neighboring cells from segmented cell mask images with uniquely labeled cells. Neighboring cells were defined using Euclidean distance thresholds of 0 pixels for direct cell contact and 10 pixels (0.5 μm) for close-proximity interactions (**Fig. 7B**). For each cell, overlapping neighboring labels within the defined distance threshold were extracted, excluding the target cell and background pixels, and matched to corresponding cluster identities using cell annotation metadata. Interaction ratios between selected cell-type pairs were calculated following the methodology described by Wang et al^48^. as the proportion of interactions involving the target cell-type pair relative to all observed cell–cell interactions. Differences in interaction ratios between progressor and non-progressor samples were assessed using the Kruskal–Wallis test.

### IMC neighborhood enrichment analysis

Neighborhood enrichment analysis was performed using Squidpy^13^ to quantify spatial interactions between IMC-defined cell populations. Cell adjacency graphs were generated using the sq.gr.spatial_neighbors function, and enrichment scores were calculated using sq.gr.nhood_enrichment based on the proximity of cell clusters within the connectivity graph. Average shortest distances between reference and target cell populations were additionally computed to characterize spatial relationships.

To account for variability in cell number across images, permutation-based normalization was applied following histoCAT^49^ framework, comparing observed interactions to randomized spatial distributions. Enrichment scores (z-scores) were extracted per image for downstream modeling and visualization. Heatmaps of average neighborhood enrichment were generated using squidpy.pl.nhood_enrichment, and in situ spatial maps were visualized using scanpy^50^ function scanpy.pl.spatial. For visualization purposes, enrichment z-scores were normalized to a range of −3 to 3.

### IMC supervised binary classification

A Random Forest classifier was trained to predict progression status using IMC-derived cellular and spatial features, including cell composition ratios, cell densities, and neighborhood enrichment scores. An initial set of 62 variables was included in the model and standardized prior to training (mean = 0, variance = 1) to ensure comparability across features. Model performance was evaluated using LOOCV to assess predictive robustness. Feature selection was performed using VSURF (Variable Selection Using Random Forest) in R to iteratively refine the input variables and identify the most informative predictors for classification. The optimized feature set was then used to train the final Random Forest model, which achieved an area under the receiver operating characteristic curve (AUC) of 0.8126. Feature importance scores were extracted from the final model to identify the variables contributing most strongly to progression prediction.

### FUME-TCRseq

FUME-TCRseq was performed as previously described^29^. Briefly, RNA extracted from FFPE tissue scrolls using the Roche HighPure FFPET RNA Isolation Kit, followed by DNase treatment, reverse transcription, and multiplex PCR amplification targeting TCRβ V regions, followed by indexed library preparation. Libraries were purified using magnetic bead cleanup, quantified by Qubit and TapeStation, pooled, and sequenced on Illumina MiniSeq or NextSeq platforms using 150 bp paired-end reads with PhiX spike-in. Sequencing reads were merged using Vsearch, and TCR clonotypes and CDR3 sequences were identified using the Decombinator pipeline. Downstream repertoire analyses were performed using Immunarch and VDJtools. TCR repertoire diversity was assessed by calculating the number of unique clonotypes per sample, both as raw counts and normalized to the specific tissue area analyzed. Clonal similarity between progressor and non-progressor samples was evaluated using the Morisita-Horn overlap index to quantify repertoire sharing.

## Supplementary Information

**Supplementary Figure 1.**
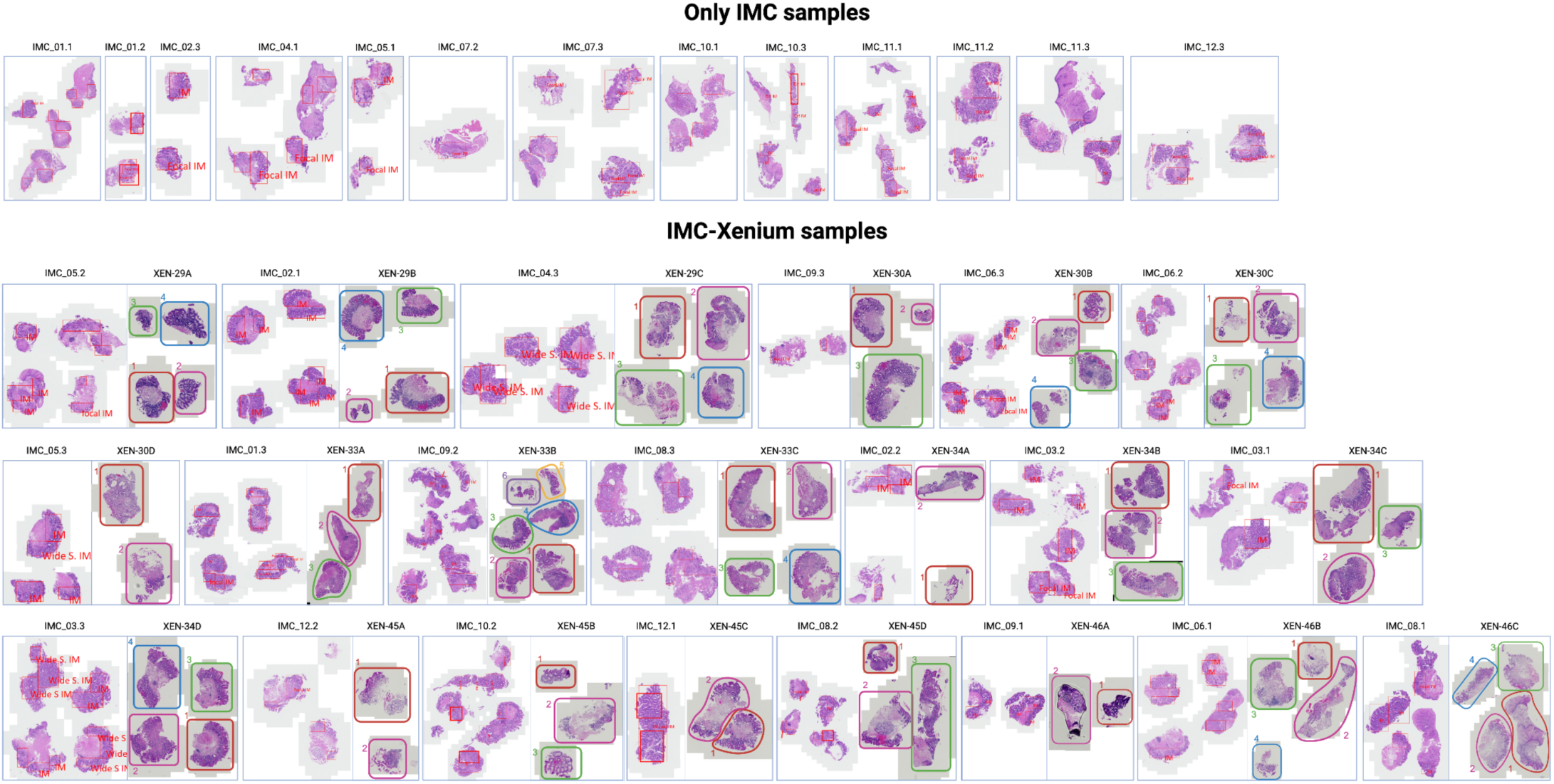
Xenium and IMC ROIs are derived from non-adjacent tissue sections. Thirteen samples are present only in IMC. Xenium profiling uses whole biopsy sections as ROIs, whereas IMC defines ROIs as subregions within the tissue.

**Supplementary Figure 2.**
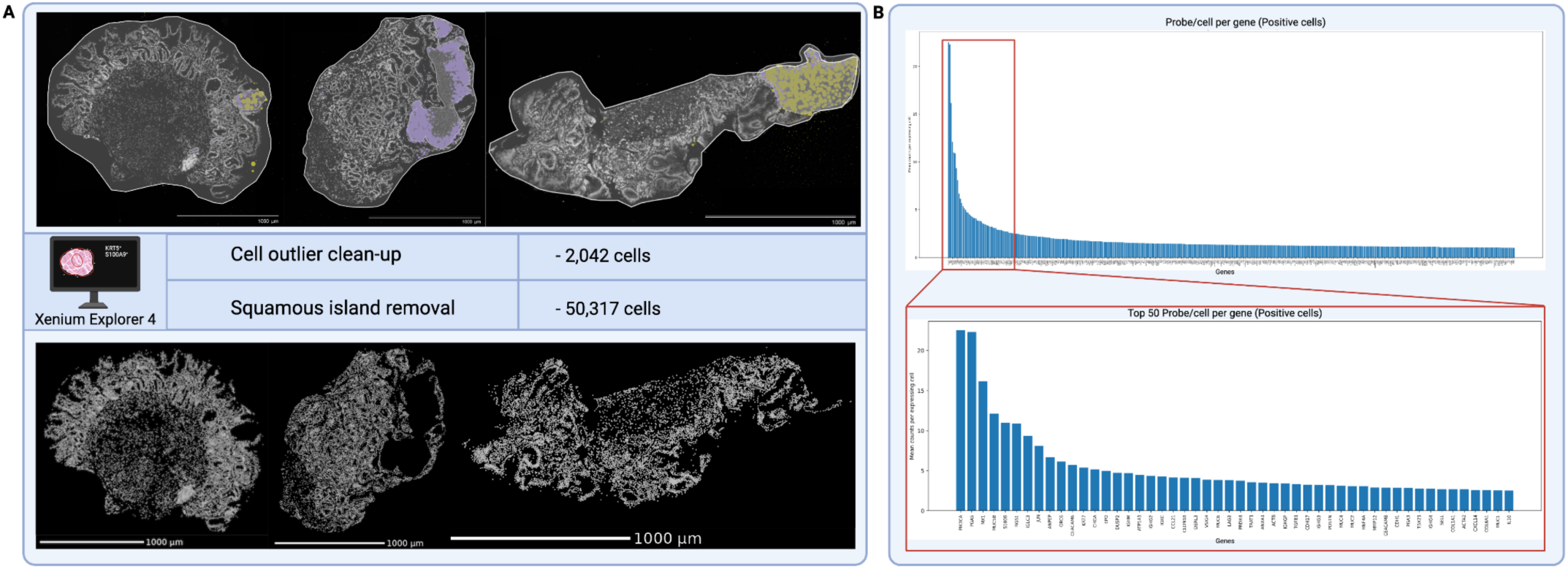
Quality control prior to clustering. **A,** Tissue mask used to remove cells outside of tissue and exclude squamous islands, as described in Methods. **B,** Bar plot showing the average probe-per-cell per gene.

**Supplementary Figure 3.**
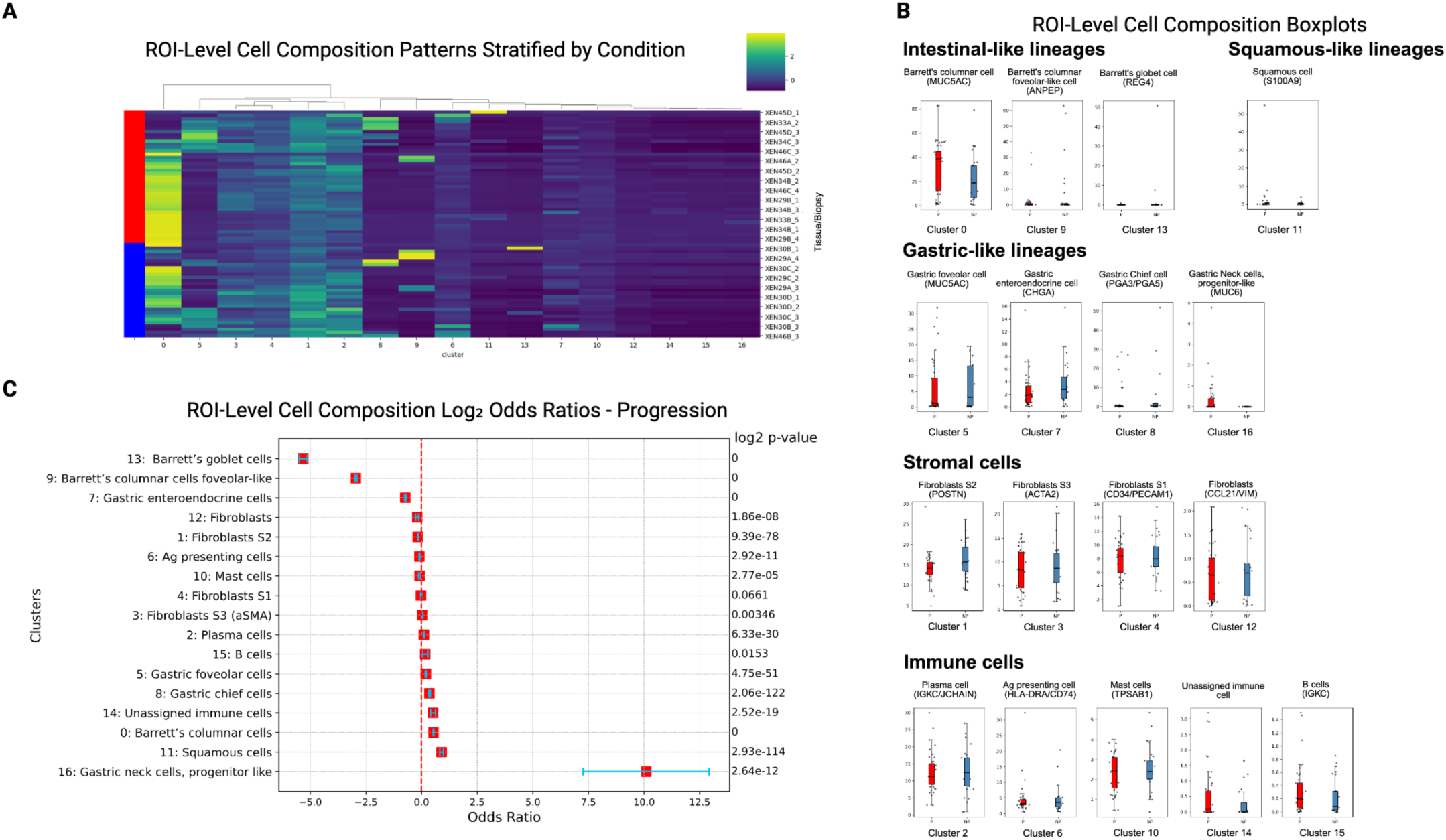
ROI-level cell composition patterns by progression status reveal limited differentiation. **A,** Heatmap of cell composition across ROIs, stratified by condition (red = Progressor, blue = Non-progressor). **B,** Forest plot of log₂ odds ratios for progression, highlighting populations enriched or depleted in Progressor ROIs. **C,** Boxplots of cell composition across annotated cell populations.

**Supplementary Figure 4.**
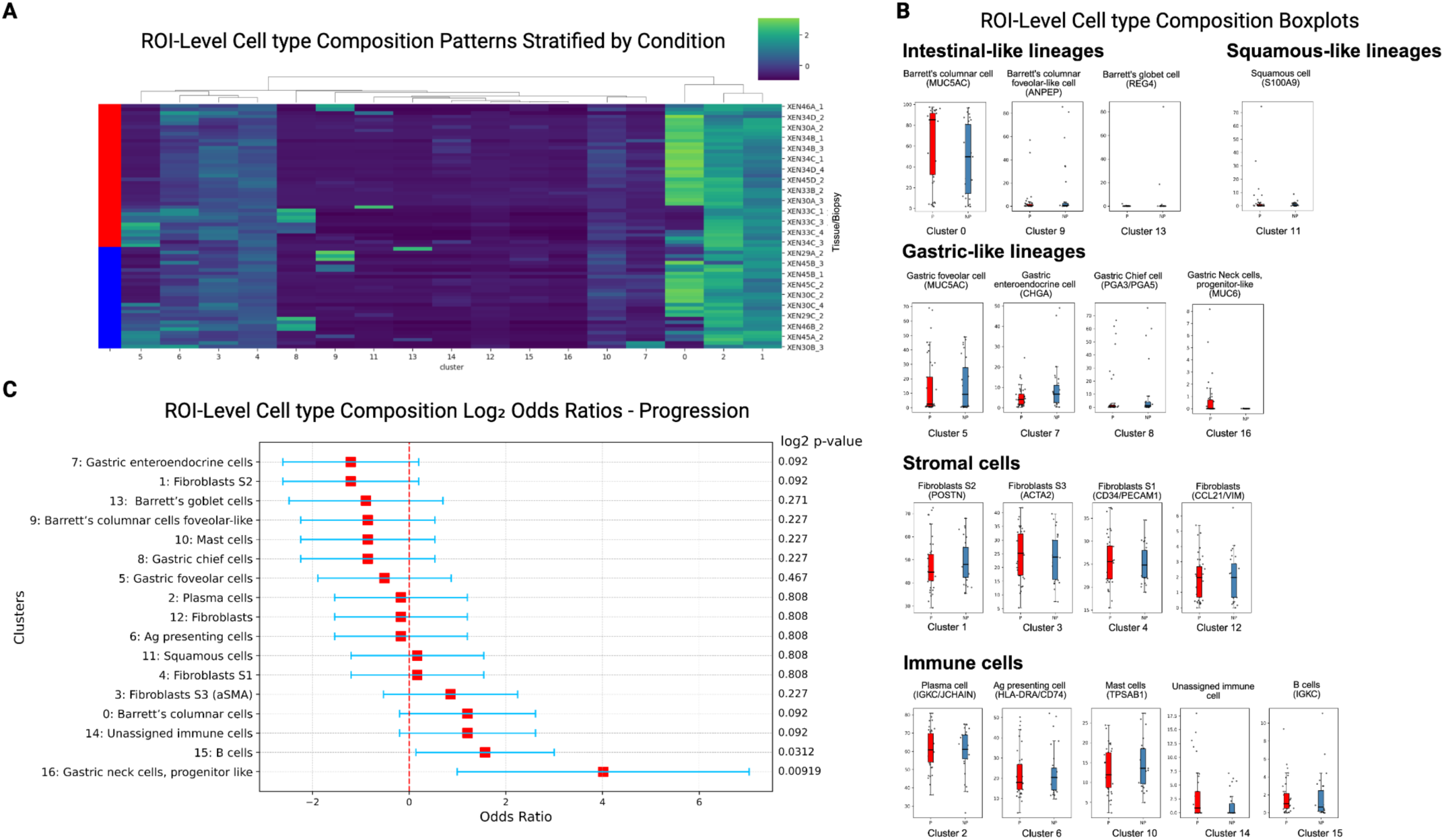
ROI-level cell composition by cell type patterns by progression status reveal limited differentiation. **A,** Heatmap of cell composition by cell type across ROIs, stratified by condition (red = Progressor, blue = Non-progressor). **B,** Forest plot of log₂ odds ratios for progression, highlighting populations enriched or depleted in Progressor ROIs.**C,** Boxplots of cell composition by cell type.

**Supplementary Figure 5.**
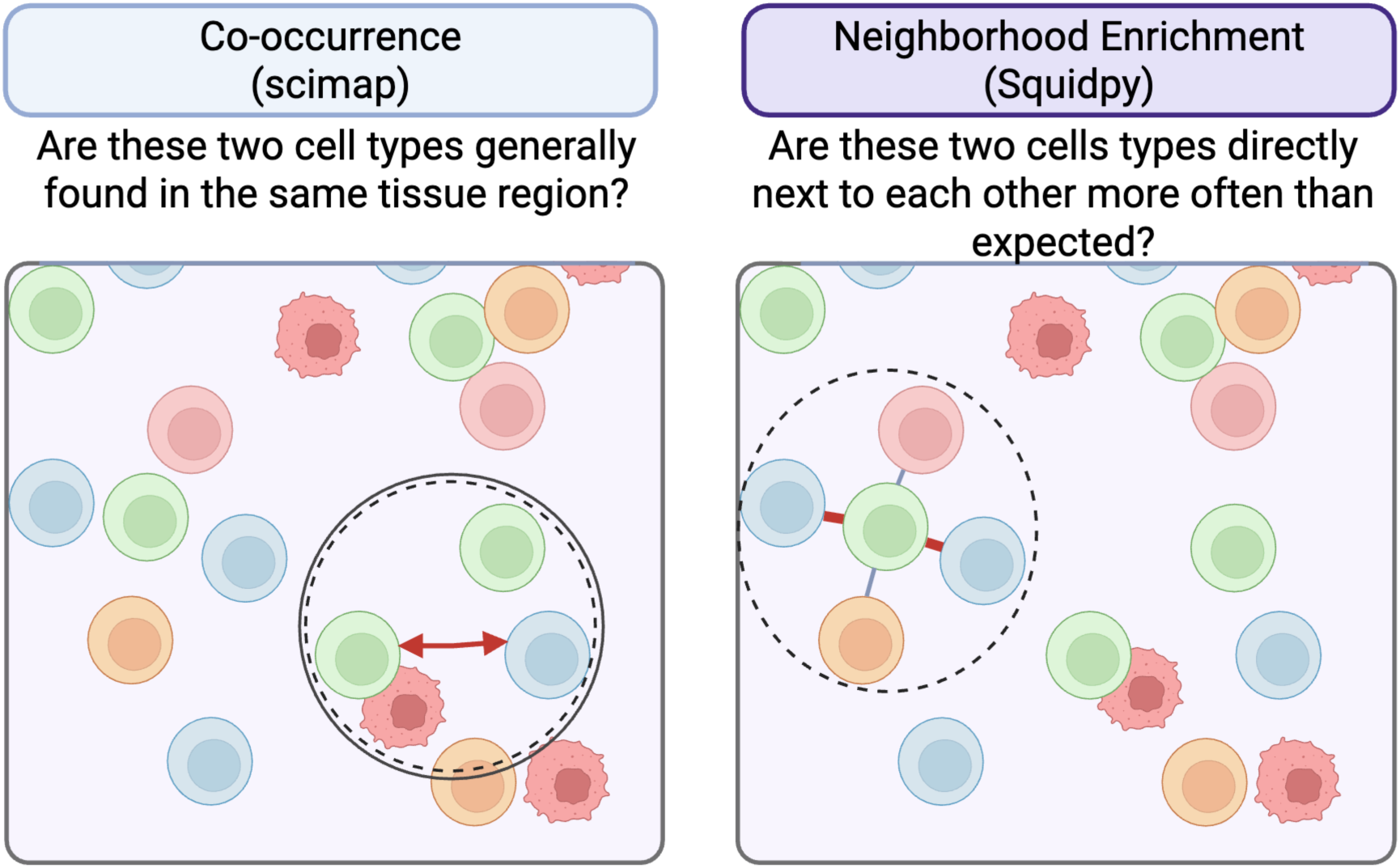
Schematic illustrating the spatial analysis framework used for cell–cell interaction analysis. Visual comparison of co-occurrence and neighborhood enrichment approaches for quantifying spatial relationships

**Supplementary Figure 6.**
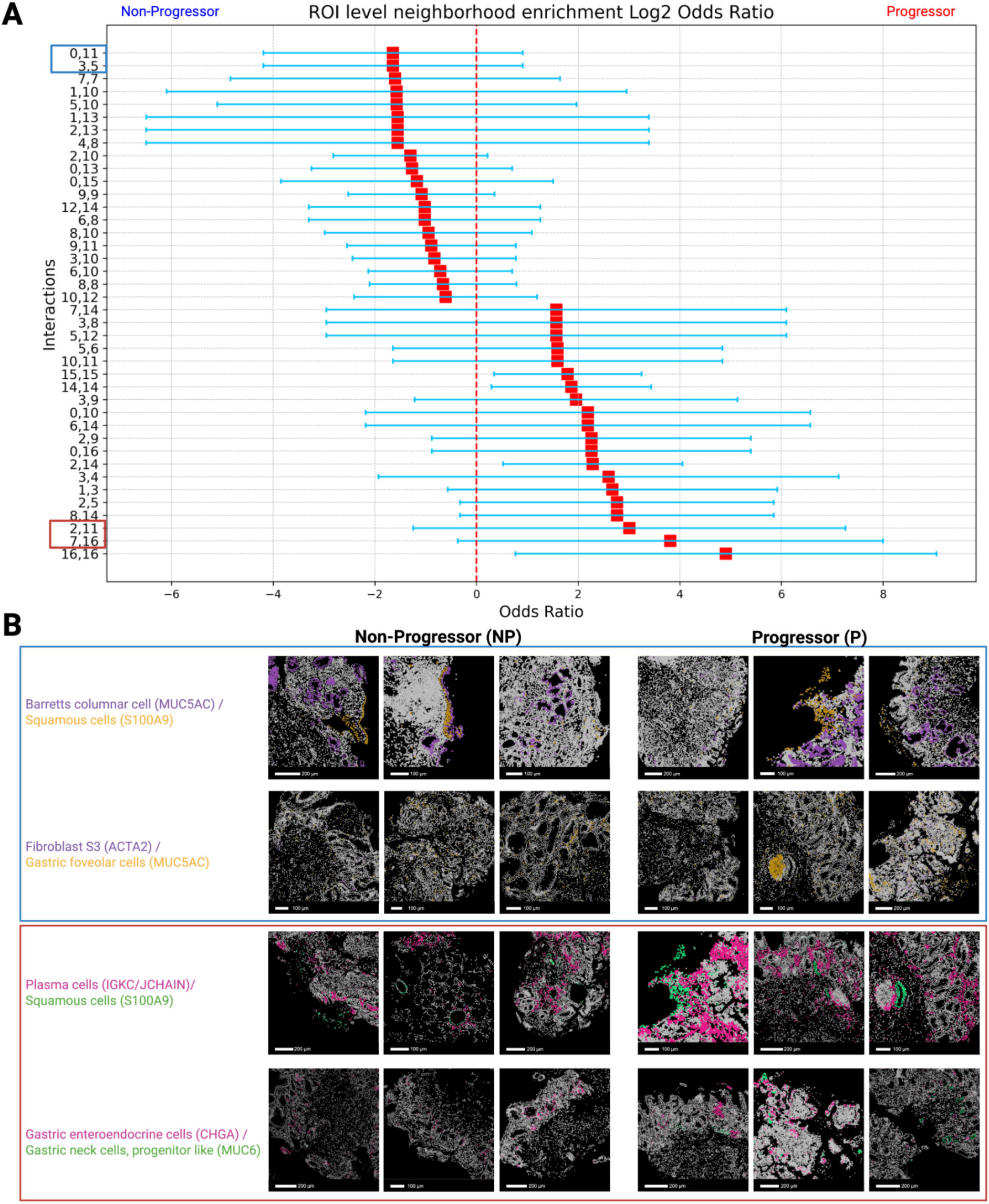
Spatial neighborhood enrichment patterns show modest differences by progression status. **A,** Forest plot of log₂ odds ratios for ROI-level neighborhood enrichment interactions, highlighting cell pairs enriched or depleted with progression. **B,** Representative spatial maps showing selected interactions in NP (blue) and P (red) ROIs, illustrating condition-specific differences in direct cell adjacency.

**Supplementary Figure 7.**
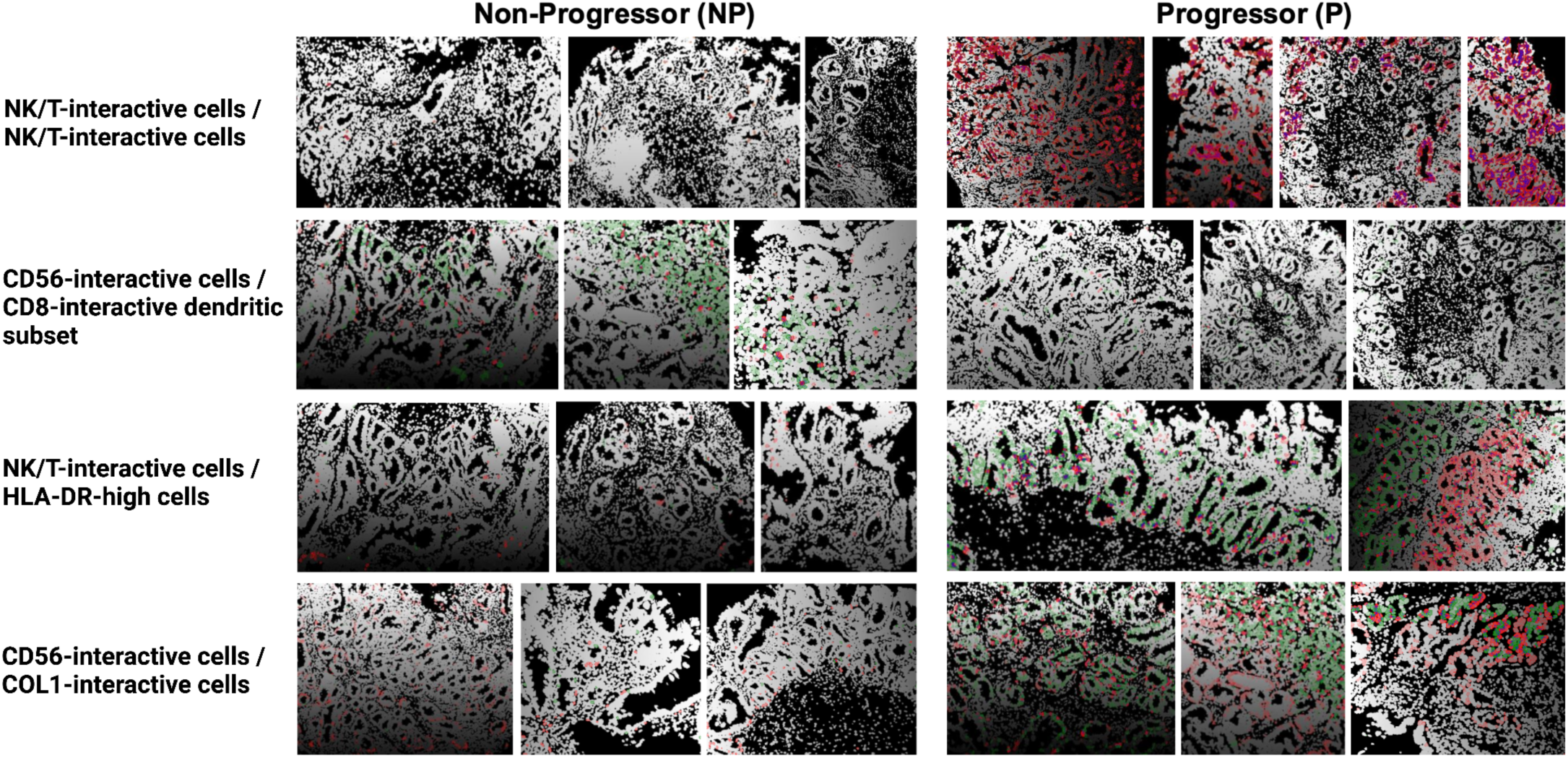
Representative spatial maps of progression-associated cell–cell interactions. Representative spatial maps of selected interactions identified by IMC-based spatial analysis in Non-progressor and Progressor ROIs.

**Supplementary Figure 8.**
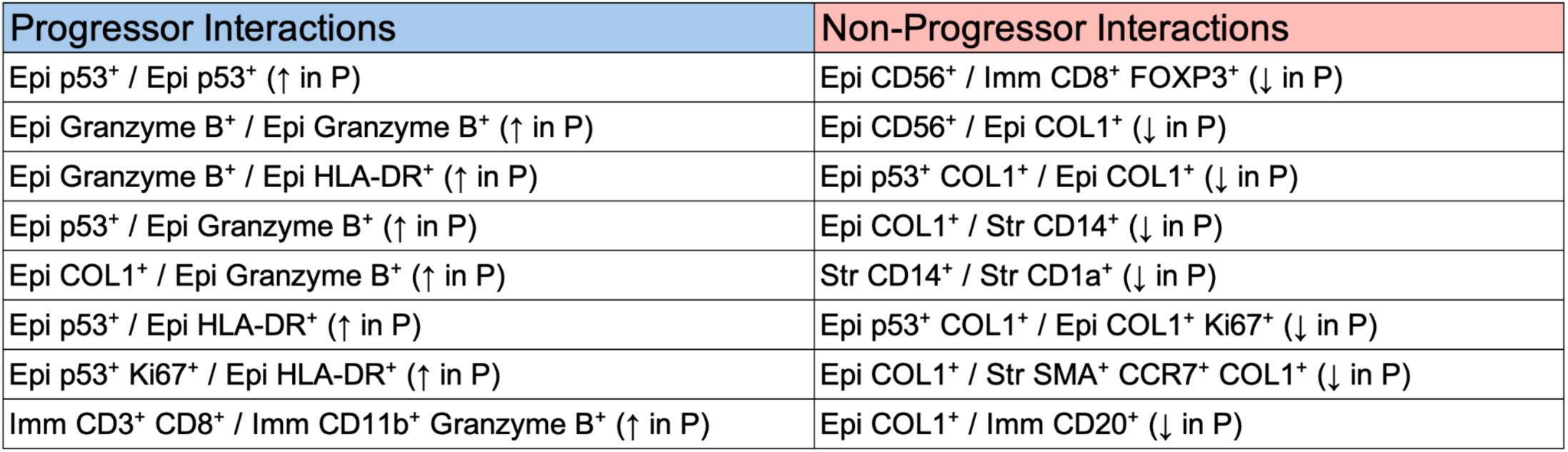
Random Forest–identified progression-associated interaction features. Cell–cell interaction features associated with progression status identified through Random Forest classification analysis.

---

**Supplementary Table 1**

Supplementary Table 1

File uploaded

**Supplementary Table 1**

Supplementary Table 2

File uploaded

